# A large-scale genome-based survey of acidophilic Bacteria suggests that genome streamlining is an adaption for life at low pH

**DOI:** 10.1101/2021.11.05.467356

**Authors:** Diego Cortez, Gonzalo Neira, Carolina González, Eva Vergara, David S. Holmes

## Abstract

Genome streamlining theory suggests that reduction of microbial genome size optimizes energy utilization in stressful environments. Although this hypothesis has been explored in several cases of low nutrient (oligotrophic) and high temperature environments, little work has been carried out on microorganisms from low pH environments and what has been reported is inconclusive. In this study, we performed a large-scale comparative genomics investigation of more than 260 bacterial high-quality genome sequences of acidophiles, together with genomes of their closest phylogenetic relatives that live at circum-neutral pH. A statistically supported correlation is reported between reduction of genome size and decreasing pH that we demonstrate is due to gene loss and reduced gene sizes. This trend is independent from other genome size constraints such as temperature and G+C content. Genome streamlining in the evolution of acidophilic Bacteria is thus supported by our results. Analyses of predicted COG categories and subcellular location predictions indicate that acidophiles have a lower representation of genes encoding extra-cellular proteins, signal transduction mechanisms and proteins with unknown function, but are enriched in inner membrane proteins, chaperones, basic metabolism, and core cellular functions. Contrary to other reports for genome streamlining, there was no significant change in paralog frequencies across pH. However, a detailed analysis of COG categories revealed a higher proportion of genes in acidophiles in the following categories: “Replication and repair”, “Amino acid transport” and “Intracellular trafficking”. This study brings increasing clarity regarding genomic adaptations of acidophiles to life at low pH while putting elements such as the reduction of average gene size under the spotlight of streamlining theory.

## 1. Introduction

Significant differences in genome sizes (number of base pairs per genome) have been detected between closely related lineages of prokaryotes isolated from a broad spectrum of environments and across multiple phylogenetic lineages, with genome sizes down to 1.2 Mb in free living Bacteria and differences of over 45% genome size between members from the same genus (Konstantinidis and Tiedje, 2004, Dufresne et. al., 2005, Lynch, 2006, Giovannoni et. al., 2014, Martinez-Cano et. al., 2015, Bentkowski et. al., 2015, Rodríguez-Gijón et. al., 2021). Small or reduced genomes, also termed streamlined genomes, have been widely observed in microorganisms adapted to live in low nutrient niches, such as cosmopolitan marine bacterioplankton (Giovannoni et. al., 2005, Schneiker et. al., 2006, Swan et. al., 2013, Luo et. al., 2014, Sun and Blanchard, 2014, Graham and Tully, 2021), rivers (Nakai et. al., 2016), slow growers in anoxic subsurfaces (Chivian et. al., 2008, McMurdie et. al., 2009), and in a wide range of extremophiles such as bacteria adapted to supersaturated silica (Saw et. al., 2008), halophiles (López-Pérez et. al.2013, Min-Juan et. al., 2016), thermophiles (Sabath et. al., 2013, Saha et. al., 2015, Gu et. al., 2020), psychrophiles (Dsouza et. al., 2014, Goordial et. al., 2016), and alkaliphiles (Suzuki et. al., 2014). Differences in genome size have been reported for aerobes versus anaerobes (Nielsen et. al., 2021) and for microorganisms living in warmer versus cooler environments (Lear et. al., 2017, Sauer and Wang, 2019) and in bacterial pathogens (Murray et. al., 2021).

Streamlining theory proposes that genome reduction is a selective process these organisms undergo that promotes their evolutionary fitness (reviewed in Giovannoni et. al., 2014). The theory suggests that a smaller genome reduces the energy cost of replication and, by encoding fewer gene products, there is a concomitant reduction of cell size that could optimize transport and nutrient acquisition (Button, 1991, Sowell et. al., 2009). Some marine microorganisms with streamlined genomes have been found to have proportionately fewer genes encoding transcriptional regulators and an overall lower abundance of mRNA transcripts per cell, potentially reducing the cost of transcription and translation (Cottrell and Kirchman, 2016). These results are congruent with the observed correlation between regulatory network complexity and genome size (Konstantinidis and Tiedje, 2004). Genome size reduction is also observed in symbiotic microorganisms (Baker et. al., 2010, Gao et. al., 2014), but it has been theorized that this phenomenon differs to the streamlining of free-living bacteria as the former lose genes by genetic drift due to function redundancy between the host and the symbiont, while the latter would lose them by intense selective pressure (McCutcheon and Moran 2012, Giovannoni et. al., 2014), although recent evidence has argued otherwise (Gu et. al., 2020).

Any organism that grows optimally at low pH can technically be classified as an acidophile. However, because there are many neutrophiles (optimum growth ∼pH 7) that successfully grow at around pH 6 or lower, it is useful from a practical point of view to define acidophiles as those microorganisms that grow optimally below pH 5 and make a distinction between moderate acidophiles that grow optimally between pH 5 and about pH 3.0 (Foster, 2004, Dopson, 2016, Benison et. al., 2021) and extreme acidophiles that grow below pH 3 (Johnson, 2007). The latter are particularly challenged for survival and growth as they face a proton concentration across their membranes of over 4 orders of magnitude (Baker-Austin and Dopson, 2007, Slonczewski et. al., 2009). Acidophilic microorganisms have been identified in all three domains of life (Johnson and Hallberg, 2003), but currently more genomic information is available for prokaryotic acidophiles (Archaea and Bacteria) (Cárdenas et. al., 2016, Neira et. al., 2020).

Our current understanding about genome streamlining in acidophiles comes from a limited number of observations. It has been reported that the genomes of several acidophilic microorganisms, such as *Methylacidiphilum, Ferrovum* and *Leptospirillum* (domain Bacteria) and *Picrophilus* (domain Archaea) are smaller (2.3, 1.9, 2.3 and 1.5 Mb, respectively) compared to their closest neutrophilic phylogenetic relatives (Angelov and Liebl, 2006, Hou et. al., 2008, Ullrich et. al., 2016, Vergara et. al., 2020). Genome reduction in acidophiles has been discussed as a mechanism to reduce energy costs to survive in extremely low pH environments where organisms must deploy multiple energy-intensive acid resistance mechanisms to maintain a circumneutral cytoplasmic pH (Hou et. al., 2008, Ullrich et. al., 2016, Zhang et. al., 2017, Vergara et. al., 2020) while thriving in often nutrient scarce and heavy metal polluted low pH environments (Johnson 1998, Dopson et. al., 2003, Johnson and Hallberg, 2008). Despite this progress, there remains much to be discovered about genome reduction in acidophiles. With the increased availability of genome sequences of acidophiles (Cárdenas et. al., 2016, Neira et. al., 2020), we shed light on whether there is a statistically supported correlation of genome reduction with low pH and, if so, what are its biological implications.

## 2. Materials and Methods

### 2.1 Data procurement and management

#### 2.1.1 Genome information

Genomes of 345 bacterial acidophiles together with their associated growth and taxonomic data were obtained from AciDB (Neira et. al., 2020). This set of genomes was modified for the present study in three ways: i) only free-living Bacteria were considered. For example, symbionts such as *Ca. Micrarchaeum* were discarded; ii) organisms without an identified phylum affiliation were also discarded and iii) seven new genomes and their associated metadata from acidophiles have been added since the publication of AciDB. This resulted in an initial dataset of 342 genomes of acidophiles. In addition, 339 genomes were collected from non-acidophiles (growth optima, pH 5-8). These included 222 genomes of neutrophiles (growth optima, pH 6-8) that were the closest phylogenetic relatives to the acidophiles as identified using NCBI taxonomy (Schoch et. al., 2020), GTDB (Chaumeil et. al., 2020) and AnnoTree (Mendler et. al., 2019), resulting in an equal taxonomic representation of genomes of acidophiles and their neutrophilic phylogenetic relatives. Genome sequences were downloaded from the National Center for Biotechnology Information (NCBI) and the Joint Genome Institute (JGI). Genomes were filtered for quality using CheckM v1.0.12 with cutoffs for completeness >80% and contamination <5% (Parks et. al., 2015). This resulted in a final data set of 597 high quality bacterial genomes, comprising 264 genomes from acidophiles (pH <5) and 333 genomes from non-acidophiles (pH 5-8). Genome information is provided in Supplementary Table 1.

Genome average nucleotide identity (ANI) was determined using fastANI v1.3 with 4 threads (Jain et. al., 2018). A cutoff of 95% average nucleotide identity was defined (Kim et. al., 2014) to group identical or highly similar genomes into species clusters. Genomic characteristics, proteomic data and associated metadata are reported as the means of each group for all plots. This reduced data bias due to over-representation of some highly sequenced species.

#### 2.1.2 Growth pH and temperature

Optimal growth pH and temperature of a species were downloaded from AciDB (Neira et al., 2020). For new species with sequenced genomes not yet deposited in AciDB, information for optimal growth pH and temperature was extracted from the literature. When no description of these optima was available, they were defined as the midpoint of the growth range reported for the strain or closely related strain as described by Neira et al., 2020. For metagenomes, the reported environmental data were used to determine optimum pH and temperature.

### 2.2 Proteome analyses

#### 2.2.1 Protein annotations

Genome annotations were downloaded from NCBI (www.ncbi.nlm.nih.gov) or JGI (img.jgi.doe.gov). Genomes without an existing annotation were annotated with prokka v1.13.3 (Seemann, 2014). A proteome table was generated for each genome, that includes information for each predicted protein, including size, predicted subcellular localization, functional annotation with COGs and Pfams, COG category, presence of signal peptide and ortholog group. Unless stated, all software was run with default options.

#### 2.2.2 Ortholog groups

To define ortholog groups, reciprocal BLASTP was performed within each genome by using all the proteins in its predicted proteome as queries against a database of the same proteins. A coverage of 50%, a sequence identity of 50% and an e-value of 10-5 were used as cutoffs (Tettellin et. al., 2005, Naz et. al., 2020). Protein pairs that follow these conditions were assigned to the same ortholog family if one or both were the best scored BLASTP hit of the other. Ortholog groups will also be referred as protein families.

#### 2.2.3 Subcellular localization

Subcellular locations were assigned to each predicted protein using PSORTb v3.0 (Yu et. al., 2010), which predicts either cytoplasmatic, inner membrane, exported, outer membrane, periplasmic for gram negative Bacteria and cell wall for gram positive Bacteria. An “unknown” tag is assigned to proteins whose subcellular location could not be predicted. This was complemented with signal peptide identification, which was assigned using SignalP v5.0b that predicts the presence of signal peptides for translocation across the plasmatic membrane by either the Sec/SPI (standard system), Sec/SPII (lipoprotein signal peptide system) or the Tat/SPI (alternative system) translocation/signal peptidases (Almagro et. al., 2019). All three positive predictions were binned together and tagged as “Has Signal Peptide”. Proteins were sorted by both subcellular localization and signal peptide presence.

#### 2.2.4 Pfam and COG functional annotations

Pfams were assigned to predicted proteins using Pfam_scan v1.6 (Finn et. al., 2016) under Pfam version 32.0 (El-Gebali et. al., 2019), which contains a total of 17929 different functional annotations including protein families and clans. An e-value of <10^−5^ was applied as a cutoff for Pfam predictions of protein function. The pfam with the lowest e-value was assigned to each protein. COG annotations were assigned with the web tool eggNOG-mapper v5.0 (Huerta-Cepas et. al., 2019) under the December 2014 version of the COG database, which contains 4632 functional annotations (Galperin et. al., 2015). The percentage of ortholog groups that have a Pfam assignment (Mistry et. al., 2021) or a COG assignment (Galperin et. al., 2021) were calculated for each proteome. The percentage of ortholog groups belonging to each COG category was also calculated. In addition, Pfam assignments were used for the analysis of intra-protein family size variation and to determine the percentage of proteins with an annotation.

#### 2.2.5 Paralog frequencies

Paralog families were defined as ortholog groups with two or more proteins from the same proteome. The percentage of proteins that belong in paralog families was calculated for each COG category in relation to the total number of proteins in the category. The same procedure was repeated for the full proteome.

### 2.3 Statistical analyses

A python script was developed to gather, filter, organize and analyze the data from the organisms’ genomes and proteomes (van Rossum, 1995). Data distributions were statistically analyzed using the following methods. The scipy library (Virtanen et. al., 2020) was used for linear fittings (with the *“linregress”* module), binomial test (with the *“stats*.*binom_test”* module) and Pearson’s linear correlation coefficient (with the *“stats*.*pearsonr”* module). A two-sided mode was used for all the tests. P-value thresholds used for statistical significance were 0.05, 0.01 and 0.001. For estimation of correlation in potentially heteroscedastic distributions, a Generalized Least Squares was applied using the module *“regression*.*linear_model*.*GLS”* within the statsmodels library (Seabold and Perktold, 2010). For multi-testing analyses, the false discovery rate (FDR) was used to determine statistical significance using the Benjamini/Hochberg procedure (Benjamini and Hochberg, 1995) with the “*stats*.*multitest*.*multipletests*” module also within the statsmodels library. A q-value of 0.05 was used for Pearson’s correlation p-values. The q-value is the upper limit of the rate of the findings (null hypothesis rejections) that is expected to be a false positive. Principal component analysis (PCA) was performed with the “*decomposition*.*PCA*” module within the sklearn library (Pedregosa et. al., 2011). The number of components for dimensionality reduction was set to 2. Data was plotted using the matplotlib library (Hunter, 2007).

## 3. Results and Discussion

### 3.1 Phylogenetic distribution and associated metadata of genomes interrogated

From the 342 publicly available genomic sequences (264 high quality plus 78 low-quality genomes) of acidophilic Bacteria, 331 genomes with well-defined taxonomies (phylum and class) were mapped on to a rooted cladogram (Figure 1). The genome sequences come from 177 species distributed in 17 classes and 8 phyla out of a total of 37 recognized bacterial phyla (55 if candidate phyla are included) (Schoch et. al., 2020) (Figure 1 and Supplementary Table 1). The acidophiles are widely distributed in the cladogram supporting the idea that acidophile lineages have emerged independently multiple times during evolution (Cárdenas et. al., 2016, González et. al., 2016, Colman et. al., 2018, Khaleque et. al., 2019, Vergara et. al., 2020).

**Figure 1.**
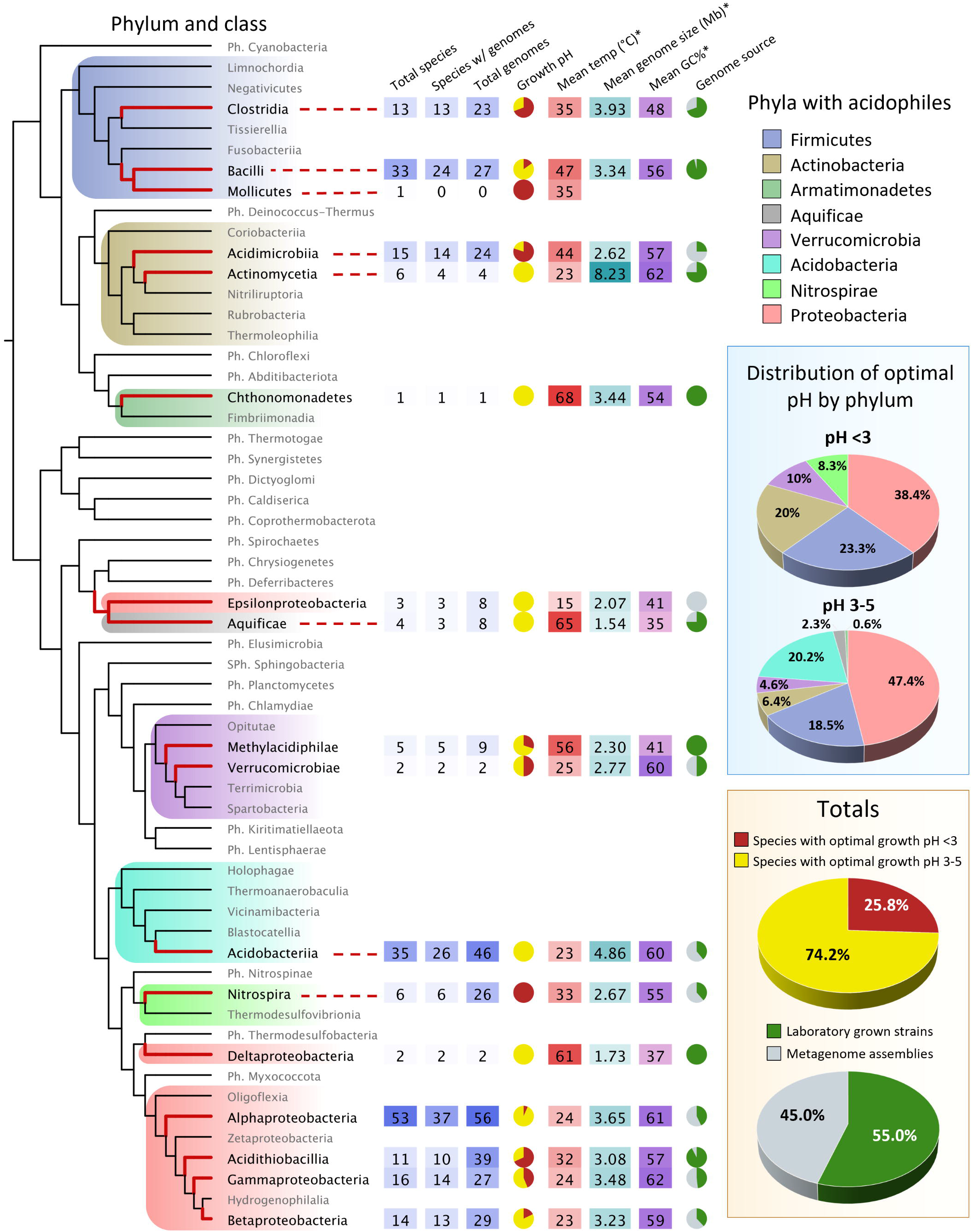
Taxonomic distribution of acidophilic genomes interrogated. A rooted cladogram displaying phyla, classes, and metadata of acidophiles with genomic data. The acidophiles are classified into those that grow optimally at pH <3 or at pH 3-5. The cladogram was constructed using AnnoTree (Mendler et. al., 2019) as a guide for phylogenetic positioning and rooted as described by Parks et. al., 2018. Phyla with acidophiles were broken down into classes. Lineages with known acidophiles are highlighted and their branches are shown with thick red lines. Dashed lines connect the acidophilic lineages with the taxon’s information when necessary. Growth pH pie charts represent the percentage of species that grow optimally at pH <3 (red) and at pH 3-5 (yellow). For both pH ranges, the percentage of acidophilic species by phyla are shown in the blue box. Genome source pie charts represent the percentage of acidophilic genomes sequenced from laboratory pure strains (dark green) versus metagenome assemblies (grey). The totals of both pie charts for all the phyla combined are shown in the yellow box. Ph. = Phylum; Sph. = Superphylum. ^*^Mean values for the acidophiles in the taxon. A more detailed table with the classes’ information can be found in Supplementary Table 2.

Figure 2 shows the distribution of acidophilic species with sequenced genomes by phylum across pH, where pH represents the optimum for growth for each species. The total number of species declines from about 60 species in the range pH 4-5 to about 10 at pH 0.5-1.5 (Figure 2A) consistent with the observation that species diversity declines in low pH environments (Bond et. al., 2000, Baker and Banfield, 2003, Johnson and Hallberg, 2003, Méndez-García et. al., 2014, Lukhele et. al., 2020, Hedrich and Schippers, 2021). These estimates are based on the distribution of acidophiles with publicly available sequenced genomes; the true richness of acidophile diversity is likely to be much higher and will probably increase as more acidic econiches are sampled using metagenomics approaches.

**Figure 2.**
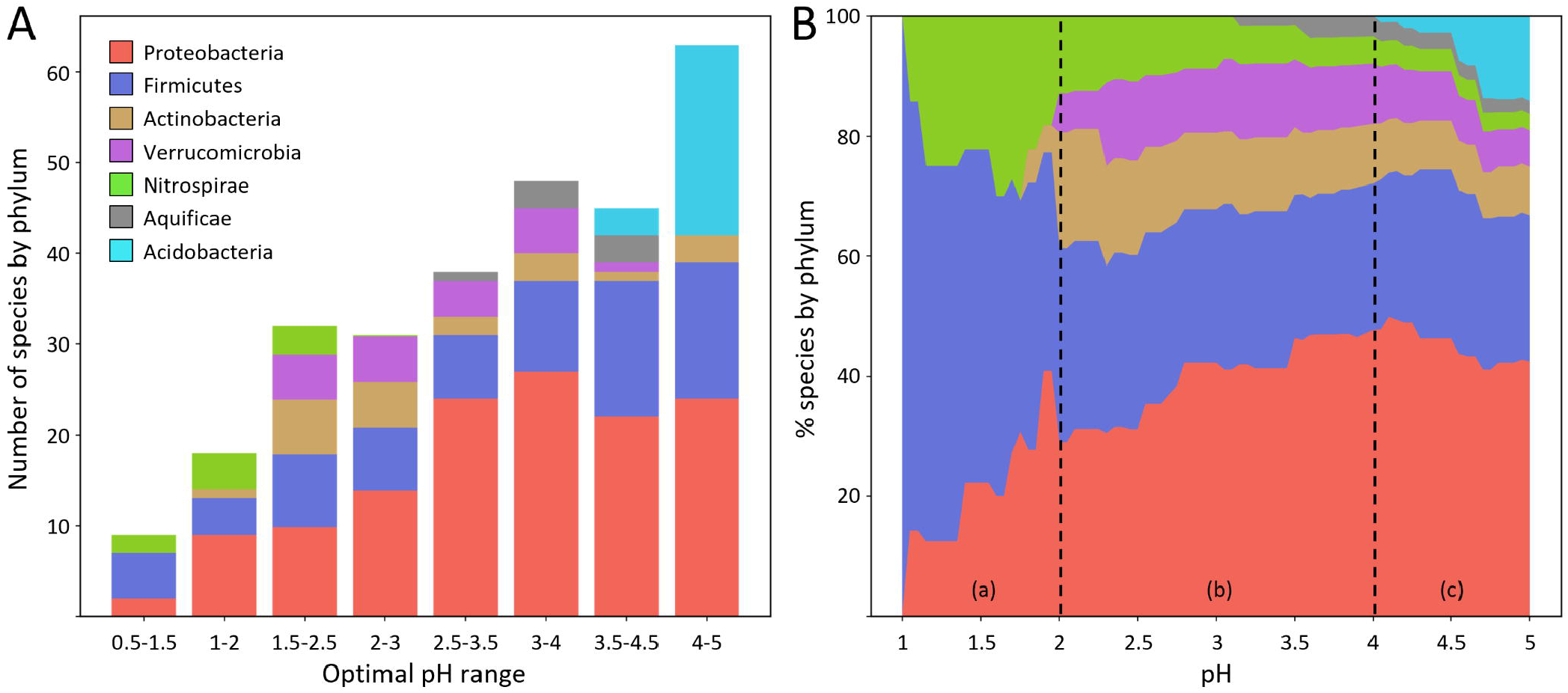
Distribution of acidophilic species with sequenced genomes by phylum across pH. Phylum *Armatimonadetes* has only one acidophilic species and is not shown. **(A)** Histogram of species number grouped by phyla across pH in overlapping increments of one pH unit. Phyla are color coded. **(B)** Cumulative plot of relative abundance (%) of acidophiles across pH. Percentages indicate species that can live at or below a given pH. Color coding of phyla is the same as A. (a), (b) and (c) indicate pH ranges 1-2, 2-4 and 4-5 respectively.

Figure 2B shows the distribution of species by percentage across pH. The results have been divided into three sections (a-c) for discussion. Section (a) with a pH range of 1.0 to 2.0 is dominated by species in the phyla Proteobacteria, Firmicutes and Nitrospirae in approximately equal proportions around pH 2 and by Firmicutes at pH 1. Section (b) shows the species distribution in the range pH 2 to 4. Acidophilic species of the phylum Proteobacteria are the most prevalent in this range but exhibit a declining percentage with decreasing pH. Species of Actinobacteria and Verrucomicrobia are represented about equally but both phyla have few representatives below pH 2. Species of Aquificae are present in a low percentage (∼ 3%) down to about pH 3, beyond which there are no representative genomes. Section (c) shows the species distribution in the range pH 4 to 5. All seven phyla (eight, if one includes the one species from Armatimonadetes) have species in this range but Acidobacteria show a declining percentage from pH 5 to pH 4 below which there are no representative genomes.

### 3.2 Genome size as a function of pH

A scatterplot of genome size across optimal growth pH shows declining genome sizes from about 4.5Mb for circum-neutrophiles to an average of about 3.4Mb for extreme acidophiles (Figure 3). There are no large genomes (>5Mb) for bacteria that grow below about pH 4, whereas large genomes including up to about 10Mb are present in acidophiles that grow between pH 4 and pH 5 and in neutrophilic relatives of the acidophiles that grow from pH 5 to pH 8. A linear regression model fitted to the data shows a tendency that is statistically significant with a positive Pearson’s correlation coefficient of 0.19 and a p-value of 2.97^*^10^−5^, implying genomes are smaller at lower pH. However, there is evidence of heteroscedasticity in the plot. We applied Generalized Least Squares Regression (GLS) to take into account heteroscedasticity, and a p-value of 1.8^*^10^−3^ was obtained supporting the proposed relationship between pH and genome size.

**Figure 3.**
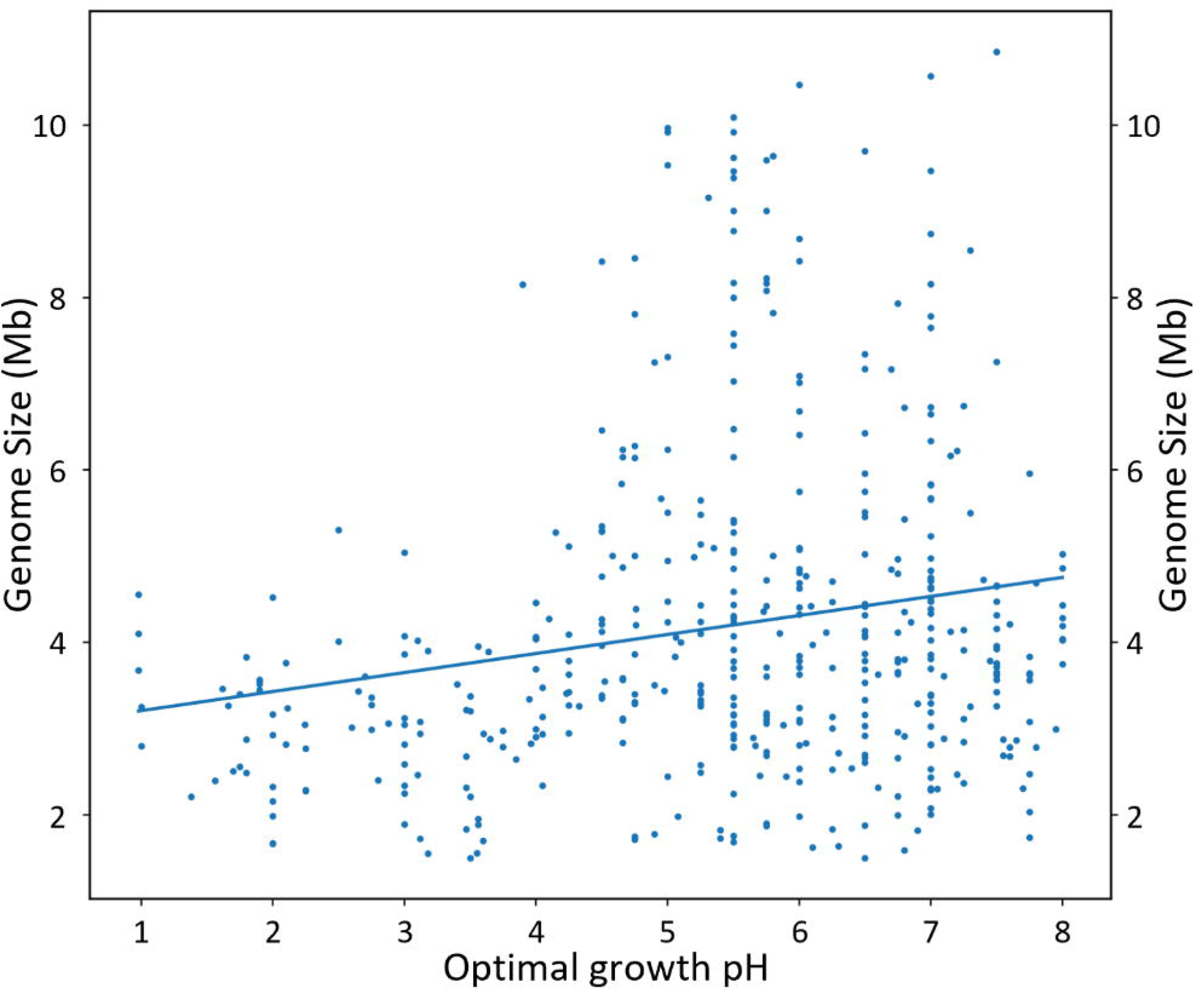
Scatterplot of genome size (Mb) of bacterial acidophiles and their most closely related extant, circum-neutral relatives versus optimal growth pH. Each point corresponds to a different species. A linear regression curve has been fitted to the data with a Pearson’s correlation coefficient of 0.19 and a p-value of 2.97^*^10^−5^. Generalized Least Squares (GLS) p-value was 1.8^*^10^−3^.

However, the presence of heteroscedasticity suggests the possibility that other variables, in addition to pH, may contribute to the determination of genome size. To address this issue, we investigated potential contributions of growth temperature and genomic G+C content on the distribution of genome size across pH. Many acidophiles are also moderate or even extreme thermophiles (Johnson and Hallberg, 2003, Capece et. al., 2013, Colman et. al., 2018) and temperature has been suggested to be a driving force for genome reduction (Sabath et. al., 2013). Genome size has also been associated with G+C content, where organisms with relatively low genomic G+C content tend to have smaller genomes (Veloso et. al., 2005, Almpanis et. al., 2018).

We evaluated how these factors are correlated with genome size and pH. Temperature is negatively correlated with genome size (Pearson’s correlation coefficient, −0.34; p-value, 2.9^*^10^−13^) (Figure 4A) and G+C is positively correlated with genome size (Pearson’s correlation coefficient, 0.48, p-value 1.9^*^10^−25^) (Figure 4C). A negative correlation between genome size and temperature has recently been reported for extreme acidophiles of the *Acidithiobacillus* genus (Sriaporn et. al., 2021). However, no statistically supported correlation is observed between temperature and pH (Pearson’s correlation coefficient, −0.01; p-value 0.84) (Figure 4B), nor between G+C content and pH (Pearson’s correlation coefficient, −0.06; p-value 0.22) (Figure 4D). Therefore, while both temperature and G+C content have a strong influence on genome size, they appear to act independently of the relationship between pH and genome size.

**Figure 4.**
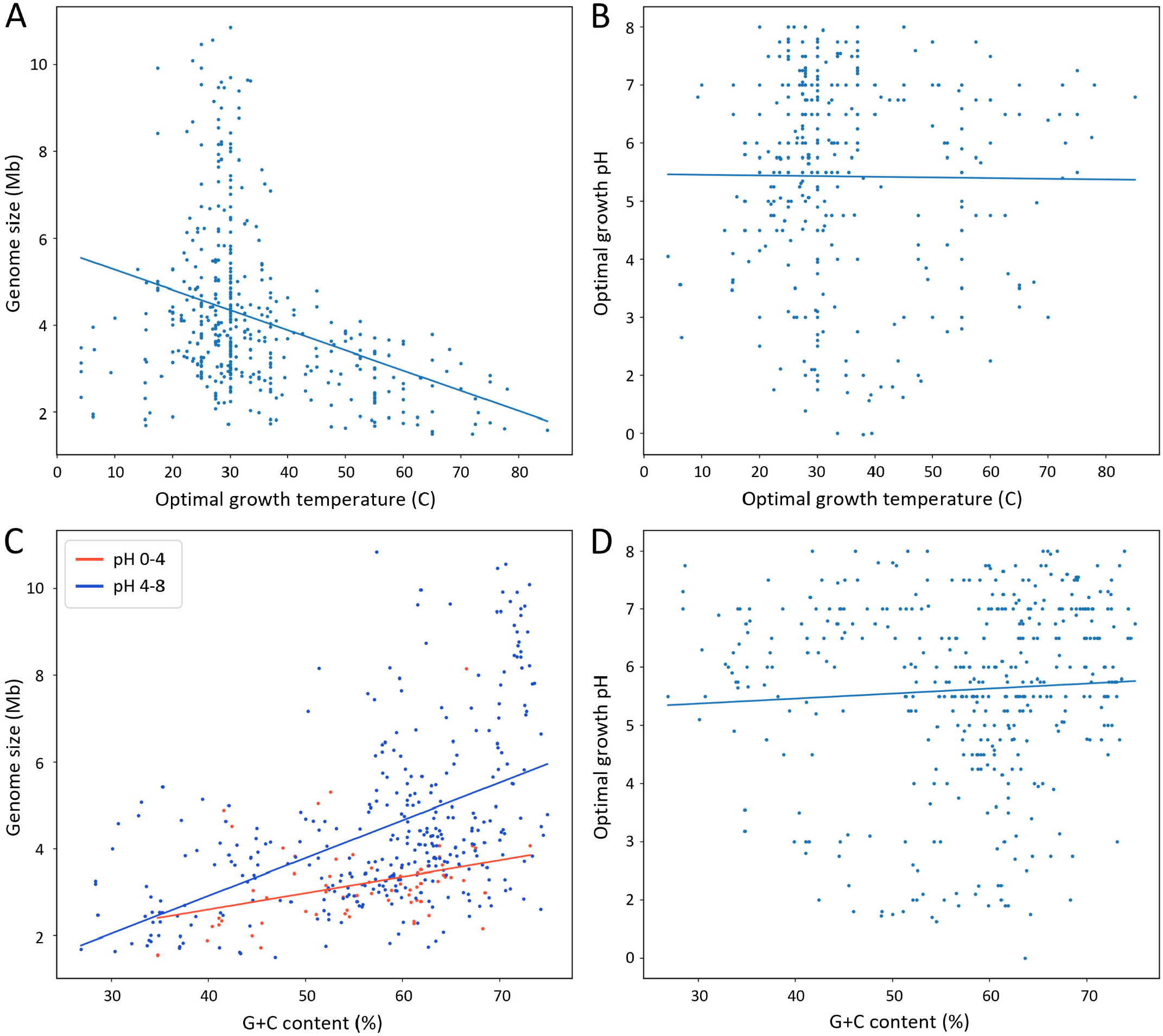
Scatterplots showing correlation of genome size and pH versus optimal growth temperature and G+C content of the species in the dataset. **(A)** Genome size vs optimal growth temperature. Pearson’s correlation coefficient is −0.34 with p-value 2.9^*^10^−13^. **(B)** Optimal growth pH versus optimal growth temperature. Pearson’s correlation coefficient is −0.01 with p-value 0.84. **(C)** Genome size versus G+C content. Here, data were separated by pH ranges. Pearson’s correlation coefficients were 0.34 and 0.50, with p-values 4.7^*^10^−3^ and 1.5^*^10^−22^ respectively for pH 0-4 and pH 4-8. The overall Pearson’s correlation coefficient and p-value were 0.48 and 1.91^*^10^−25^, respectively. **(D)** Optimal growth pH versus G+C content. Pearson’s correlation coefficient is −0.06 with p-value 0.22.

To investigate further the interplay of pH, temperature and G+C content with genome size, we performed dimensionality reduction and visualization via principal component analysis (PCA) (Jolliffe, 2005). As seen in Figure 5, the directions of the loading vectors show temperature is negatively correlated with both G+C content and genome size, while genome size is positively correlated with both G+C content and pH. This is also depicted in how the smallest genomes are found in thermophiles (optimal temperature >55°C, rightmost cluster) followed by extreme acidophiles (optimal pH <3, upmost cluster), while the biggest genomes are found in a high G+C content group (leftmost cluster). Conversely, the orthogonality of the loading vectors suggests no correlation is observed between pH and temperature or between pH and G+C content. Therefore, when considering all variables at once, the same results are observed as when the variables were individually assessed (Figure 4), providing additional evidence that neither G+C content nor temperature affect the correlation between pH and genome size, rather multiple driving forces can independently exert their influence on genome size.

**Figure 5.**
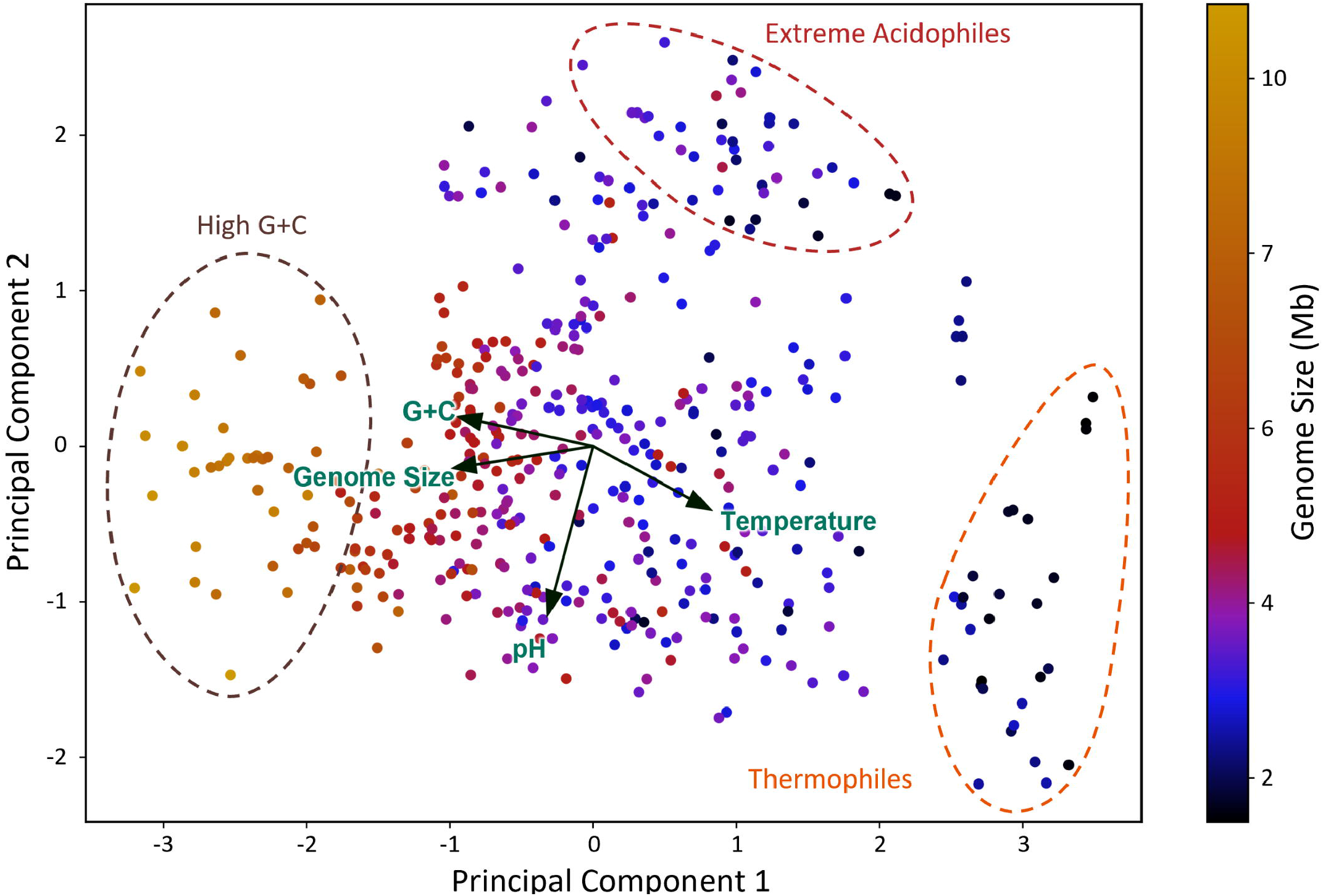
Principal component analysis of multiple variables potentially influencing genome size. Dimensionality reduction was performed by PCA, inputting the optimal growth pH, optimal growth temperature, G+C content and genome size of each species in the dataset. A biplot was constructed showing the loadings of each variable as arrows at the center of the plot and the distribution of the principal components. The average genome size of each species is shown as a color scale. Three clusters within the dotted circles are highlighted for their distinctives features.

### 3.3 Genetic mechanisms involved in genome size changes

#### 3.3.1 Hypothetical schema

Given the observation that genome size is negatively correlated with pH in acidophiles, we aimed to determine what genomic processes influence this relationship. Figure 6 shows a diagrammatic representation of genetic mechanisms that have been postulated to be involved in genome expansion or reduction in Bacteria and Archaea (Keeling and Slamovits, 2005, Sabath et. al., 2013, Giovannoni et. al., 2014, Gillings, 2017, Kirchberger et. al., 2020, Rodríguez-Gijón et. al., 2021, Westoby et. al., 2021). Genome size changes could result from having (i) changes in number of orthologous families (A, Figure 6) or paralogous genes (B, Figure 6); (ii) genome compaction/expansion resulting from changes in the number of intergenic nucleotides including alteration in the frequency of overlapping genes (C, Figure 6) (reviewed in Kirchberger et. al., 2020) and (iii) smaller or larger genes, including loss/gain of domains (D, Figure 6).

**Figure 6.**
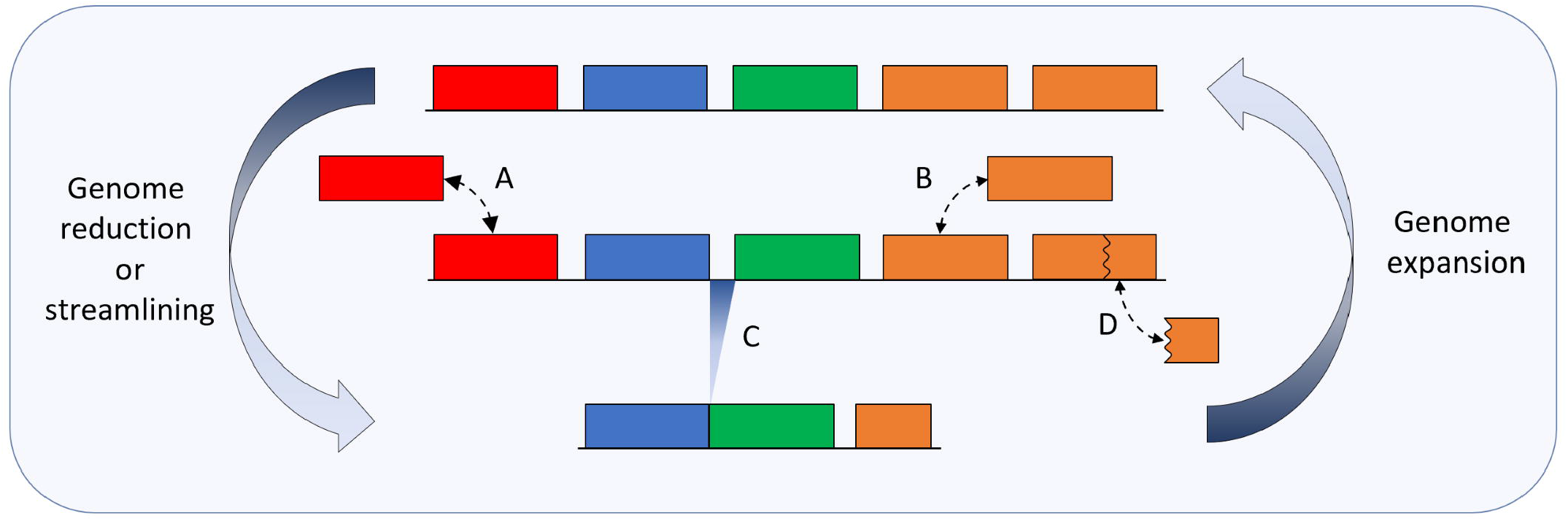
Diagrammatic representation of genetic mechanisms involved in genome size changes. **Top row**, five genes of a hypothetical genome. Orange boxes indicate paralogous genes. **Middle row**, processes involved in genome size changes where A and B represent gene loss/gain of single copy genes or paralogous genes respectively, C shows intergenic space reduction or expansion, which we refer to as genome compaction, and D shows gene size reduction or increase. **Bottom row** reduced or streamlined genome relative to the starting genome shown in top row; alternatively, the starting genome before expansion to genome shown in top row. Large blue arrows indicate time or direction of evolutionary events. Small dotted bidirectional arrows show hypothetical insertion or deletion events.

Based on the schema shown in Figure 6, we investigated the contribution of the different mechanisms in genome size changes in acidophiles across pH. Annotated open reading frames (ORFs) were used as surrogates for “genes”. A caveat is that ORF prediction depends on the quality of the genome sequence, where poor quality genomes frequently have incorrectly annotated chimeric and truncated ORFs that confound subsequent identification of genes (Klassen and Currie, 2013). We minimized these potential errors by analyzing only genomes that had passed a high quality CheckM filter (Parks et. al., 2015). However, even high-quality genomes are prone to errors of ORF annotation especially in the identification of correct translation start sites (Korandla et. al., 2020) which will impact predictions of gene and intergenic spacer sizes. Currently, there are no computational program for ORF prediction that is flawless, including GenBank (Korandla et. al., 2020), and we expect that future work will improve the annotations of ORFs used in our study.

#### 3.3.2 Reduction/expansion of gene (ORF) number

The number of protein coding genes (ORFs) of each genome under interrogation was plotted as a function of optimal growth pH of the species. The results indicate that there is a statistically significant reduction (Pearson’s coef.: 0.18; P-value: 1.25^*^10^−4^) of the average number of ORFs per organism across pH from an average of about 4100 ORFs/organism at pH 7 to about 3200 ORFs/organism at pH 2 (Figure 7A). This has been regarded as possibly the most predominant mechanism for genome size changes (Konstantinidis and Tiedje, 2004) and this is likely also true for our dataset (Supplementary Figure 1).

**Figure 7.**
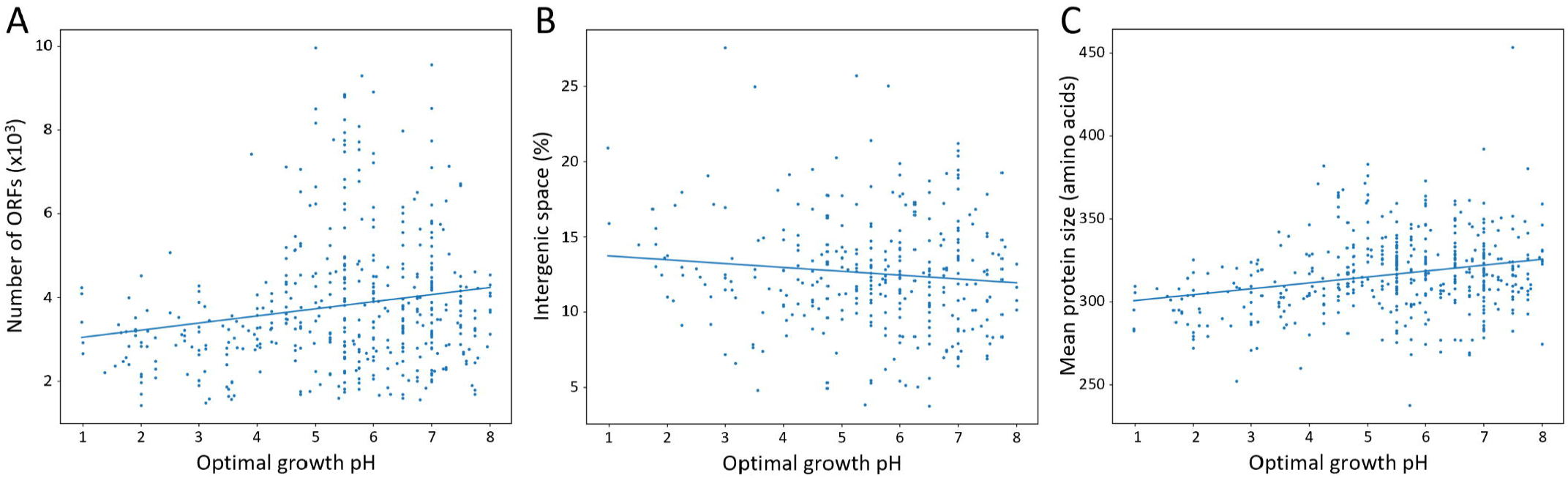
Factors influencing genome size of acidophiles across optimal growth pH. Every point corresponds to the average for a different species. **(A)** Number of genes (ORFs, open reading frames) across pH. Pearson’s correlation coefficient is 0.18 with p-value 1.25^*^10^−4^. **(B)** Intergenic space vs pH. Intergenic space is defined as genome size minus the sum of the nucleotide length of all protein coding genes as defined by ORFs of a genome divided by genome size, in percentage. A stricter genome quality filter of 97% completeness and 2% contamination was used in this analysis to minimize missannotation errors due to fragmented genomes. Pearson’s correlation coefficient is −0.11 with p-value 0.06. **(C)** Average ORF length per genome across pH. Pearson’s correlation coefficient is 0.25 with p-value 4.03^*^10^−8^.

#### 3.3.3 Reduction of intergenic spacers as a possible contributor to genome compactness

It is well established that bacteria have compact genomes with an average protein-coding density of 87 % with a typical range of 85–90 % (McCutcheon and Moran 2012). Genome size reduction could occur by decreasing the amount of DNA occupied by intergenic spacers, for example by promoting the frequency of overlapping genes (Veloso et. al., 2005, Saha et. al., 2015, Kreitmeier et. al., 2021). This strategy has been especially exploited in compacting viral genomes (Pavesi, 2021).

To evaluate whether a reduction in the fraction of the genome dedicated to non-protein coding DNA contributed to smaller genomes observed in acidophiles, we calculated the percentage of intergenic spaces (IG) dedicated to the total genome content across pH. IG was calculated as genome size (bp) - ∑ bps of all ORFs in a genome, expressed as a percentage of the total bps in the genome. A smaller % IG implies greater genome compaction. A tendency was observed for % IG to increase as pH growth optima declines (Figure 7B), however, this trend is not statistically significant (Pearson’s coef. = −0.11, p-value 0.06). A potential problem in the interpretation of this result stems from uncertainties in the identification of ORFs, most notably by errors in the identification of the correct site of initiation of protein coding regions (Korandla et. al., 2020). This influences the estimation of the percentage of intergenic genomic DNA.

#### 3.3.4 Reduction/increase of gene (ORF) size

The average size of ORFs (as the number of amino acids of the predicted proteins) per genome was plotted as a function of pH (Figure 7C). There is a statistically supported positive correlation (p-value 4.03^*^10^−8^) between average ORF size and pH, with an average size of 320 amino acids at pH 7 to 300 at pH 2. This indicates acidophiles have shorter proteins in average, which could be produced by a loss of larger proteins or by gene size reduction (Figure 6, mechanism D) or possibly both.

To quantify gene size reduction in acidophiles, we analyzed the protein sizes of several conserved Pfams in the dataset (Figure 8). We observed that the conserved Pfams with reduced protein sizes in acidophiles are over 5 times as many as the conserved Pfams with increased sizes (Figure 8 A, binomial test p-value 2.1^*^10^−13^). This result accounts mainly for changes in the predominant domain architectures, implying these proteins in acidophiles likely have fewer domains. For example, the biotin requiring enzyme was mainly found in single domain proteins below pH 5, while in neutrophiles it can often be found next to other domains such as the dihydrolipoamide acyltransferase (Supplementary Table 3). This inclination towards protein size reduction is also observed in a collection of conserved Pfams that are also in single copy and predominantly in single domain architectures (Figure 8 B, binomial test p-value 7.4^*^10^−3^). This result accounts mainly for loop size reductions and domain size reductions. Such is the case of the ribosomal protein L19 that in acidophiles lacks long loops and is 4 amino acids shorter on average (Supplementary Table 4).

**Figure 8.**
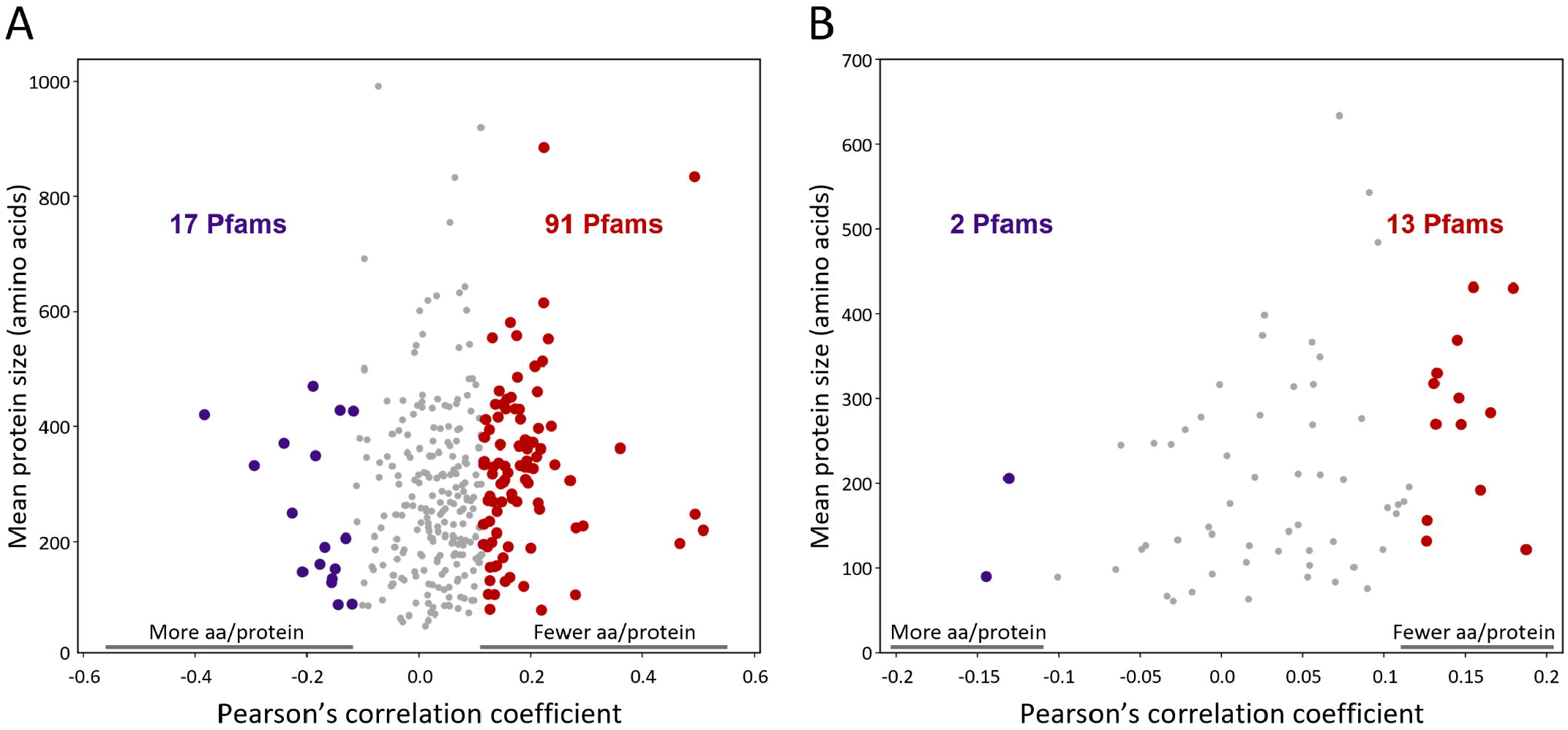
Protein size versus pH correlations for conserved Pfams. **(A)** Pfams present in over 90% of species and in a pH span of at least 6 pH units were selected for analysis. For each Pfam, the Pearson’s correlation coefficient for protein size vs organism optimal growth pH was calculated, using the species averages as data. Each point corresponds to a different Pfam. Positive correlations (91 red points to the right) indicate Pfams whose proteins are shorter at low pH while negative correlations (17 purple points to the left) are Pfams whose proteins are larger at low pH. The 25 Pfams with the lowest p-values are listed in Supplementary Table 3. **(B)** Analog to **(A**), but for a list of Pfams that in addition to being present in over 90% of the species and in a span of at least 6 pH units were also in a unique copy in the genomes (proteins with the Pfam per genome <1.1) and only one domain architecture was dominant in the proteins. These Pfams are listed in Supplementary table 4. For both plots, an FDR q-value of 0.05 was used for statistical significance. Significant correlations are shown as big points which are red for positive correlations and purple for negative correlations. Non-significant correlations are shown as small grey points.

### 3.4 Gene representativity across pH

Having established that there is a statistically supported negative correlation between genome size and optimal pH for growth and that gene gain and loss events likely contributed to this correlation, we investigated in more detail what types of genes were involved these events.

#### 3.4.1 Changes in ortholog groups representativity in acidophiles

To gain insight into the contribution of gains or losses of genes in the observed genome size changes of acidophiles (mechanism A, Figure 6), we first clustered the genes into ortholog families and systematically classified the predicted proteomes of each genome by (i) subcellular location and (ii) functional category as predicted by Pfam annotations (Mistry et. al., 2021) and COG categories (Galperin et. al., 2015). Subsequently, we mapped the frequencies of ortholog families of these categories in the genomes across pH.

##### 3.4.1.1 Changes in ortholog frequencies by sub-cellular location

Figure 9 shows the frequency of occurrence of protein families with sub-cellular location and/or signal peptide predictions expressed as a percentage of the total protein families per genome. The frequency of proteins predicted to be in the cytoplasm does not change across pH (blue data points and line, Figure 9). However, there is a statistically significant decrease (Pearson’s correlation coefficient 0.22, p-value 1.4^*^10^−6^) in the frequency of proteins predicted to have a signal peptide with decreasing pH (red data points and line, Figure 9) and a statistically significant increase (Pearson’s correlation coefficient −0.19, p-value 4.4^*^10^−5^) in the frequency of proteins predicted to be in the inner membrane with decreasing pH (orange data points and line, Figure 9). There is a small, but nevertheless statistically significant decrease (Pearson’s correlation coefficient 0.21, p-value 7.5^*^10^−6^) in the frequency of proteins predicted to be in the category “periplasm, outer membrane, cell wall and exported” with decreasing pH (green data points and line, Figure 9).

**Figure 9.**
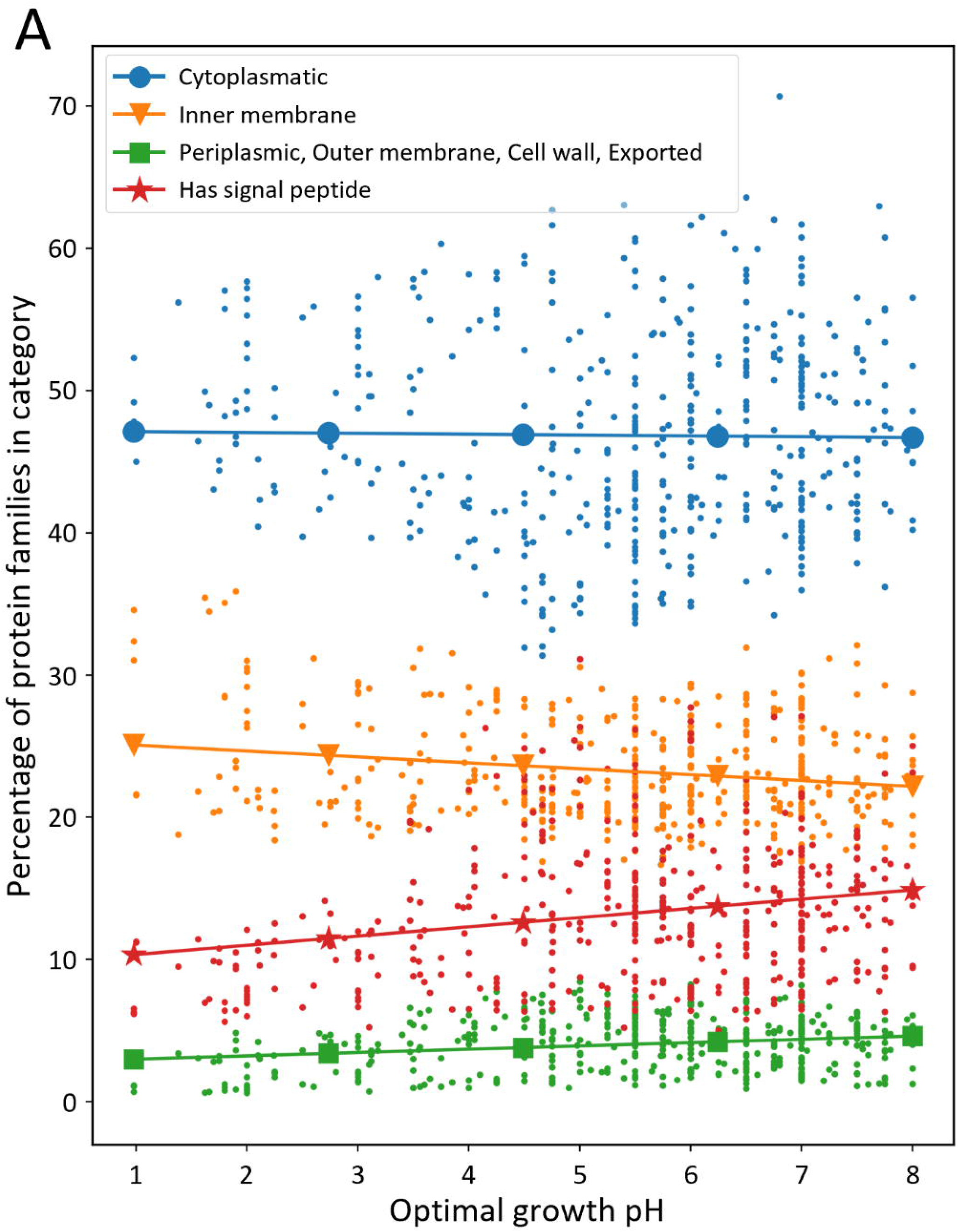
Subcellular localization and signal peptide presence of protein families across pH. PSORTb and SignalP were used to predict subcellular location of proteins and signal peptide, respectively. Each point corresponds to a species, and either subcellular localization or signal peptide presence are expressed in terms of percentage of the protein families (ortholog groups). Linear regression curves have been plotted for each category. Pearson’s correlation coefficient and p-value respectively are −0.01 and 0.77 for cytoplasmic, −0.19 and 4.4^*^10□^5^ for inner membrane, 0.21 and 7.5^*^10□ ^6^ for Periplasmic, Outer membrane, Cell wall and Exported, and 0.22 with 1.4^*^10□^6^ for proteins with a signal peptide.

The decrease in proportion of proteins with signal peptides at low pH (Figure 9) is consistent with the observation that there are correspondingly fewer proteins predicted in the category “periplasm, outer membrane, cell wall and exported” at low pH since most of these proteins require a signal peptide export mechanism to pass through the periplasmic membrane (Green and Mecsas 2016). We hypothesize that the decrease in relative frequency of proteins found outside the inner membrane in acidophiles could be due to physico-chemical challenges that such proteins would encounter as they are exposed to high concentrations of protons at low pH, potentially limiting the diversity of proteins that have evolved to survive such challenges (D’Abusco et. al., 2005, Chi et. al., 2007, Duarte et. al., 2009, 2011, Panja et. al., 2020, Chowhan et. al., 2021). We speculate that the observed enrichment of protein families predicted to be in the inner membrane in acidophiles (Figure 9) reflects the importance of such proteins in acid stress management (Lund et. al., 2014, Zhang et. Al., 2016, Vergara et. al., 2020, Hu et. al., 2020).

##### 3.4.1.2 Changes in ortholog frequencies by functional category

The contribution of gene gain or loss to genome size changes across pH were also analyzed using gene functional classification using COG and Pfam annotations. 25 functional categories are recognized in the 2014 COG database (Galperin et. al., 2015) and Pfam v32.0 contains a total of 17,929 families (El-Gebali et. al., 2019, https://pfam.xfam.org). The combination of COG and Pfam analyses provides deep and accurate coverage for searching for predicted protein function in our dataset. Figure 10 shows that the percentage of proteins per genome with a COG or Pfam annotation decreases at lower pH with statistical significance (Pearson’s correlation coefficients 0.24 and 0.14, p-values 2^*^10^−7^ and 2.6^*^10^−3^), which is not observed for small neutrophilic genomes (Supplementary Figure 3). This indicates that acidophiles have a higher proportion of putative protein coding genes that are not recognized by neither COG nor Pfam. These proteins can be classified as non-conserved, hypothetical proteins with no functional prediction, which do not have protein clusters with sufficient entries to have their own functional annotation in the COG or Pfam databases. It is possible that some of these represent poorly annotated sequences and pseudogenes. However, an intriguing possibility is that some could correspond to *bona fide* protein coding genes that are enriched in acidophiles. Their analysis could potentially yield clues about novel acid-tolerance mechanisms and other functions enriched in acidophiles. Examples of such proteins have recently been detected, although their function remain unknown (González et. al., 2016, Vergara et. al., 2020).

**Figure 10.**
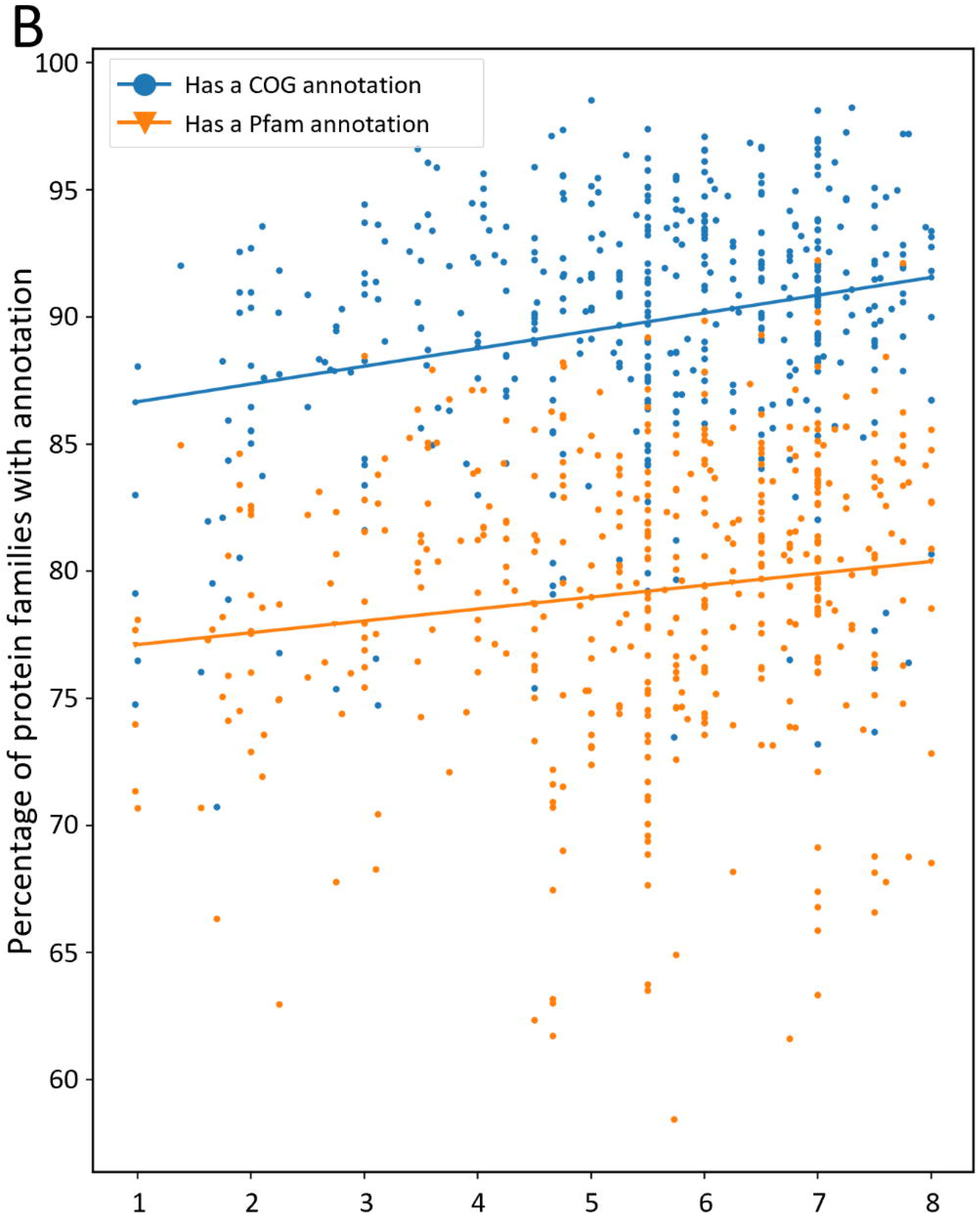
Percentage of protein families with functional classification across pH. Each point corresponds to a species. Blue data points and the blue line correspond to proteins with a COG annotation and orange data points and the orange line correspond to proteins with a Pfam annotation. Pearson’s correlation coefficients and p-values are respectively 0.24 and 2^*^10□^7^for proteins with a COG annotation, and 0.14 with 2.6^*^10□^3^ for proteins with a Pfam annotation.

An analysis of the distribution of functional categories across pH using COGs shows that acidophiles are enriched in several functions that could possibly be attributed to their distinctive metabolisms and environmental challenges (Table 1). For example, enrichment in COG L (replication, recombination, and repair) and COG O (Chaperone, post-translational modification) might reflect their need for DNA repair and protein refolding when confronted by potentially damaging stresses such as low pH, high metal concentrations and oxidative stress (Crossman et. al., 2004, Baker-Austin and Dopson, 2007, Cárdenas et. al., 2012, Dopson and Holmes, 2014). The increase in frequency of COGs C, F and H (Energy production and transport; nucleotide metabolism and transport and coenzyme metabolism and transport, respectively) could reflect enzyme and pathway requirements associated with obligate autotrophic metabolism that has been found in many acidophiles (Johnson, 1998, Johnson and Hallberg 2008). As for COG J, it is possible that as ribosomal proteins are very conserved across prokaryotic life (Lecompte et. al., 2002), they are less likely to be discarded. Future research could investigate what are the functions in this category overrepresented in acidophiles.

**Table 1.** Genomic representativity of protein families by function as defined by COG categories in acidophile genomes.

| COG Category | Pearson's correlation coefficient | p-value |
| --- | --- | --- |
| **Increased representativity in acidophiles (p-value<0.01)** |  |  |
| (L) Replication, recombination, and repair | -0.25 | 3.6*10<sup>-8</sup> |
| (F) Nucleotide metabolism and transport | -0.21 | 5.4*10<sup>-6</sup> |
| (C) Energy production and conversion | -0.21 | 8.0*10<sup>-6</sup> |
| (H) Coenzyme metabolism and transport | -0.19 | 3.0*10<sup>-5</sup> |
| (D) Cell cycle control and cell division | -0.16 | 5.2*10<sup>-4</sup> |
| (J) Translation and ribosome | -0.15 | 1.1*10<sup>-3</sup> |
| (O) Chaperones, post-translational mod. | -0.13 | 6.3*10<sup>-3</sup> |
| **Decreased representativity in acidophiles (p-value<0.01)** |  |  |
| (S) Function unknown | 0.30 | 1.3*10<sup>-10</sup> |
| (T) Signal transduction mechanisms | 0.26 | 3.4*10<sup>-8</sup> |

On the contrary, genomes of acidophiles are depleted in COG T (Signal transduction mechanisms). A depletion of signal transduction mechanisms has been observed in some marine microbes especially those that are slow growing (Gifford et. al., 2013, Cottrell and Kirchman, 2016), in the streamlined genome of the extreme acidophile *Methylacidiphilum infernorum* (Hou et. al., 2008) and in metagenomic profiling data of acidic environments (Chen et. al., 2015). The abundancy of signal transduction mechanisms generally declines with decreasing genome size, as it has been found that the number of one and two component signal transduction systems is proportional to the square of the genome size (Konstantinidis and Tiedje, 2004, Ulrich et. al., 2005, Galperin, 2005). Extensive research has been conducted on the different signal pathways and regulatory networks of acidophiles (Rzhepishevska et. al., 2007, Shmaryahu et. al., 2009, Moinier et. al., 2017, Díaz et. al., 2018, Osorio et. al., 2019). However, additional research is needed to uncover what signal pathways are not present in these organisms. Acidophiles possess several features which may explain their underrepresentation in proteins from this category, such as having small genomes, and having relatively slow growth speeds (Fang et. al., 2006, Mykytczuk et. al., 2010).

The genomes of acidophiles also have a proportionately reduced number of COG S (unknown function). These are proteins with unknown function that are conserved across multiple species and so are distinct from the category described above (Figure 10) that are not conserved across multiple species. As both are proteins with no known function, the representativity of unknown function proteins remains relatively constant across pH, but a greater number of these proteins are in multiple species in neutrophiles. It is possible that many functions assigned to COG S are found principally in neutrophilic heterotrophs whose genome sequences are the most prevalent in databases (extrapolated from the limited number of genomic sequences of acidophiles, Neira et. al., 2020) and therefore can potentially dominate the COG database.

#### 3.4.2 Paralog frequency across pH

We next examined whether the gain or loss of paralogs contributed to genome size changes (mechanism B, Figure 6). In contrast to what has been described above concerning gain or loss of specific COG and Pfam gene functions, here we explored how genome size could be influenced by the expansion or contraction of the number of genes in such families. Gene duplication, followed by functional diversification has been invoked as a major contributor to gene evolution (reviewed in Innan and Kondrashov, 2010 and Copley, 2020) and gene paralogs can be present as a significant proportion of a genome (Swan et. al., 2013). An increase in the number of paralogous protein copies (including in- and out-paralogs and xenologs, Remm et. al., 2001, Darby et. al., 2017) has been observed to be correlated with a better performance in a specific function, such as heavy metal resistance or adaptation to other multiple stressors (Kondratyeva et. al., 1995, Dulmage et. al., 2018). Relatively high paralog frequencies for proteins linked to acid resistance mechanisms have been detected in acidophiles (Ullrich et. al., 2016, Vergara et. al., 2020).

We analyzed paralog frequency changes in genomes across pH by COG categories. The COG annotation has been proved useful for gene enrichment analyses across several genomes (Galperin et. al., 2021). As seen in Figure 11 and Supplementary Figure 5, acidophiles have relatively high paralog frequencies in the COG categories “Replication, repair and recombination”, “Intracellular trafficking and secretion” and “Energy production and conversion”, but low frequencies in the COG categories “Signal transduction”, “Translation and ribosome” and “Amino acid metabolism”, as shown by statistically significant correlations (p-value <0.01). Some of the results are in concordance with the protein family representativity results (Table 1) which increases the importance of the putative contribution of these functions on acidophilic survival and adaptation.

**Figure 11.**
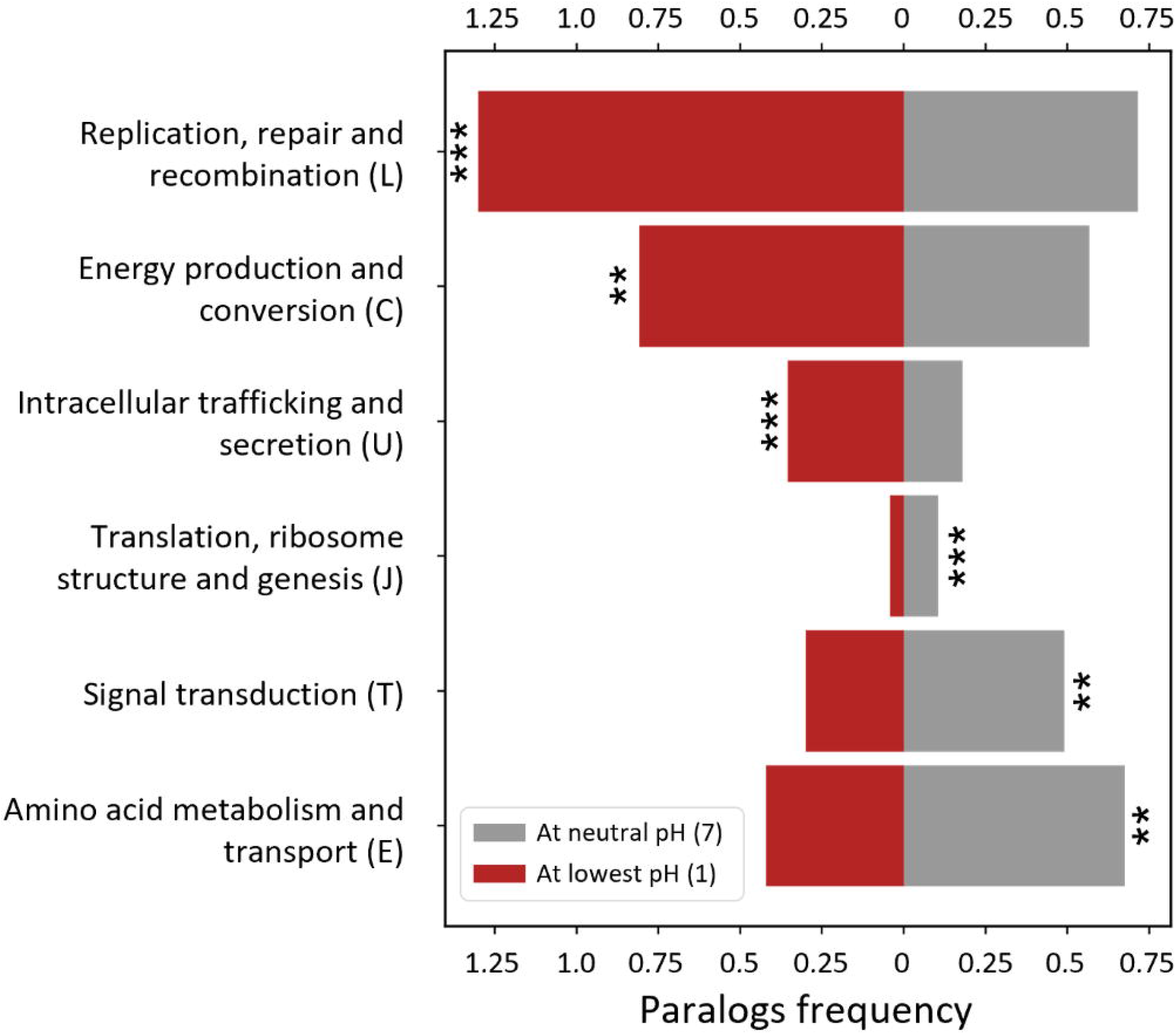
Paralog frequency vs pH by COG category. The percentage of genes (relative to the proteome size) belonging to paralog families (paralog frequency) were calculated for each COG category. Categories where the paralog frequency had a statistically significant correlation with pH (p-value <0.01) are shown. The mean duplication frequencies at pH 1 and 7 are displayed, calculated with linear regression (Supplementary Figure 5). ^**^ p-value<0.01, ^***^ p-value<0.001.

High paralog frequencies in the “Replication, repair and recombination” category in acidophiles might be attributed to a large number of transposases and integrases. The high prevalence of mobile elements in acidophilic genomes has been previously pointed out as a key factor for acidophilic evolution (Aliaga et. al., 2009, Navarro et. al., 2013, Acuña et. al., 2013, Ullrich et. al., 2016, Zhang et. al., 2017, Colman et. al., 2018, Vergara et. al., 2020). As discussed in the previous section (Table 1), DNA repair proteins might also be in several copies. These have been found to protect against oxidative stress and heavy metal stress, which acidophiles are exposed to in higher levels (Crossman et. al., 2004, Baker-Austin and Dopson, 2007, Cárdenas et. al., 2012).

The increased number of paralogous proteins from the “Intracellular trafficking and secretion” category in the acidophile genomes could result from an abundance of type II secretory systems involved in conjugation. It has been observed that these systems are frequently associated with mobile elements and are found to be particularly abundant in the flexible genomes of acidophiles (Acuña et. al., 2013, Beard et. al., 2021), suggesting that they are shared between organisms in a common econiche. In addition, vesicle related proteins might also be duplicated in acidophilic genomes, as studies show that vesicular transport (whose related functions belong in this category) is linked to biofilm formation (Jan, 2017), which in turn has been widely observed in acidophiles (Baker-Austin et. al., 2010, González et. al., 2013, Díaz et. al., 2018, Vargas-Straube et. al., 2020).

Similarly to the results of genome representativity (Table 1), the increased paralog frequencies of proteins from the “Energy production and conversion” category in acidophiles, might be related with their overrepresentation of chemolithotrophic metabolism. Some of the enzymes involved in iron or sulfur oxidation belong to this category, such as the cytochrome C, heterodisulfide reductase and quinone related proteins (Quatrini et. al., 2009, Zhan et. al., 2019). Additionally, several proteins in this category are involved in proton exporting functions, such as the H+-ATPase and the overall electron transfer chain proteins such as the ubiquinone oxidoreductase (Walker, 1992, Fütterer et. al., 2004, Feng et. al., 2015). This indicates that some genes in this category might be in high copy numbers to increase the acid resistance of acidophiles. Alternatively, it could be a consequence of the high energy requirements of maintaining a neutral internal pH (Baker-Austin and Dopson, 2007, Slonczewski et. al., 2009).

The reduced paralog frequencies in the “Signal transduction” category are concordant with their reduced genome representativity in acidophiles, and thus might be accounted by the same phenomena as previously exposed (Table 1).

The reduced number of paralogs in acidophiles in COG E “Amino acid transport and metabolism”, might be accounted for by a reduction in the number of amino acid importers that are not common in acidophiles. The predominancy of autotrophic metabolism in acidophiles could result in an inclination for these organisms towards biosynthesis of amino acids rather than uptake by active transporters. Additionally, uptake of amino acids could be harmful to acidophiles as organic acids carry protons into the cytoplasm of these organisms, short circuiting acid resistance mechanisms (Kishimoto et. al., 1990, Lehtovirta-Morley et. al., 2014, Carere et. al., 2021). The current hypothesis is that organic acids are protonated in the extremely acid medium where acidophiles grow (pH <3) becoming non-ionic and soluble in bacterial membranes, permitting diffusion into the cytoplasm (pH around 7) where they uncouple from the proton. A similar phenomenon could occur with amino acids but involving membrane transporters, as amino acids are unlikely to diffuse passively through the membrane.

As for COG J “Translation and ribosome”, their reduced paralog frequency is opposite to the increased representativity of protein families from this category in the genomes of acidophiles (Table 1). In other words, acidophiles tend to discard (or not evolve) duplicated genes from this category rather than losing core functions by relinquishing unique protein families. Further exploration is needed to determine what are the changes this category in acidophiles.

Concordantly, as there was an equilibrium between COG categories with increased and decreased paralog frequencies in acidophiles, the overall paralog frequency had no statistically significant correlation with optimal pH and remained at a relatively constant 8% average, ranging from 2% to 20% (Supplementary Figure 4). These relatively low percentages indicate that paralog frequencies are only a minor contributor to genome size changes in our dataset. Still, the constant paralog frequency across pH contradicts what has been found for other streamlined organisms, which have relatively low number of paralogs (Giovannoni et. al., 2005, Swan et. al., 2013). This unusual finding could be partially a consequence of acid resistance genes in multiple copies that would compensate the evolutionary pressure of discarding paralogs.

## 4. Additional Discussion

We have shown acidophilic Bacteria possess several streamlining elements, such as having smaller genomes, fewer ORFs and an underrepresentation of signal transduction proteins (Gifford et. al., 2013, Giovannoni et. al., 2014, Cottrell and Kirchman, 2016). However, there are several streamlining elements that we could not identify in acidophiles, such as having lower intergenic space percentages, lower paralog frequencies and proportionately fewer pseudogenes (Giovannoni et. al., 2005, Swan et. al., 2013). This could be partially attributed to the high prevalence of HGT and recombination elements in acidophiles (Aliaga et. al., 2009, Navarro et. al., 2013, Acuña et. al., 2013, Ullrich et. al., 2016, Zhang et. al., 2017, Colman et. al., 2018, Vergara et. al., 2020). A high recombination activity is prone to increase the abundancy of pseudogenes present in a genome (Holt et. al., 2009, Tutar, 2012) and could cause the observed high paralog frequencies in the Cog category L “Replication, recombination and repair”, which in turn increases the overall paralog frequencies of acidophiles. This is supported by the low paralog frequencies in COG category J “Translation and Ribosome”, which are amongst the most conserved proteins (Lecompte et. al., 2002) and thus could be an index of general paralog frequency tendencies. Additionally, streamlining as a phenomenon has been mainly described for extremely small genomes (<2Mb). While genomes as small as 1.7Mb exist in our dataset, most of the genomes are between 2-4 Mb, which could explain the absence of some streamlining elements in acidophiles.

What is observed for acidophiles then appears to differ from the classic examples of extremely streamlined organisms. However, as opposed to statistical analyses of multiple acidophilic clades, most of the studies that defined genome streamlining traits focus on a single clade and reflect on the underlying ecological variable to which attribute its genome reduction (Dufresne et. al., 2005, Giovannoni et. al., 2005, Chivian et. al., 2008, Sowell et. al., 2009, López-Pérez et. al., 2013, Luo et. al., 2014, Sun and Blanchard, 2014, Nakai et. al., 2016, Cottrell and Kirchman, 2016, Graham and Tully, 2021). The divergence in the observations from this study and others could be attributable to such difference, as single clade studies do not consider counter examples such as *Rhodococcus erythropolis*, an extreme oligotroph with a genome of over 7 Mb (Yano et. al., 2016, Retamal-Morales et. al., 2018). Nevertheless, streamlining in the evolution of acidophiles appears to be a less robust phenomenon than in thermophiles when comparing to other multi-clade statistical studies (Sabath et. al., 2013). This was also observed in our study, as shown by the stronger correlation between genome size and temperature (Figure 4A) than with pH (Figure 3) and the positioning of the lowest genome sizes in the PCA plot (Figure 5).

In terms of physiology, acidophiles possess several characteristics of streamlined Bacteria, such as relatively small cell sizes (Clark and Norris, 1996) and high generation times (Kishimoto and Tano, 1987, Fang et. al., 2006, Mykytczuk et. al., 2010). Chemolithoautotrophic metabolism is widespread amongst acidophiles (Johnson and Hallberg, 2008), which could be a bias in our study as the reduced genomes of acidophiles might be related to this overrepresentation of chemolithoautotrophs. However, some of the smallest genomes in free-living prokaryotes are heterotrophs (Giovannoni et. al., 2005, 2014) and are smaller than some of the smallest known genomes of chemolithoautotrophic prokaryotes besides methylotrophs (Raven et. al., 2013). Therefore, this is unlikely to be a major issue.

In agreement with what has been observed in Archaea (Colman et. al., 2018), the bacterial acidophiles are all nested within higher order neutrophilic lineages and no examples are observed of regression of acidophile lineages to neutrophiles, suggesting that the evolution of acidophilia is unidirectional. However, the current taxonomic distribution of acidophilic genomes is possibly affected by sampling bias, as acidic mine drainages are one of the most studied acidic environments (Johnson and Hallberg, 2003, Sharma et. al., 2016) which possibly produces an overrepresentation of organisms from these environments in the databases. Advances in metagenomics should attenuate this issue by increasing the genomic information from less studied acidophilic econiches, such as deep-sea vents (Simmons and Norris, 2002, Reysenbach et. al., 2006) and to a lesser extent solfataric fields (Itoh et. al., 2011). Possibly, entirely novel acidophilic lineages from different phyla could be discovered.

Some of the genomic traits observed in acidophiles have not been described as general features of streamlined organisms, such as lower average protein sizes and higher representativity of inner membrane proteins. These features could be novel characteristics of streamlined organisms or perhaps are specific for acidophilic adaptation. The increased representativity of inner membrane proteins is likely to be specific for acidophiles, as no statistically supported correlation was found between the representativity of these proteins and genome size in neutrophiles (Supplementary Figure 2). This is also likely true for the lower representativity of proteins found outside the inner membrane of acidophiles. In contrast, average protein size has been analyzed in previous streamlining studies on adaptation to high temperatures (Sabath et. al., 2013). A decrease in average protein size was reported for thermophiles, and a conclusion regarding thermostability adaptations (Thompson and Eisenberg, 1999, Chakravarty and Varadarajan, 2000) was reached. However, protein size changes might be a major contributor to genome size changes besides gene gain or loss. Our discovery of a decrease in average protein size in acidophiles expands the possibility beyond thermophiles that protein size reduction might be a more general mechanism for genome streamlining in stressful environments. Further research on this feature is necessary to determine whether other streamlined organisms have smaller proteins than their counterparts. Nevertheless, smaller proteins in acidophiles could also be attributable to protein stability adaptations, such as the shorter loops observed for some proteins in the inner membrane of acidophiles (Duarte et. al., 2009, 2011). The investigation of which specific protein size changes or domain rearrangements might be attributable to a survival mechanism in acidic econiches is a potential topic for future research.

Acidophiles pay the energetic toll of maintaining a proton gradient of several orders of magnitude across the inner membrane (Baker-Austin and Dopson, 2007, Slonczewski et. al., 2009). This, while proliferating in often nutrient scarce environments with multiple stressors (Johnson, 1998, Dopson et. al., 2003, Johnson and Hallberg, 2008). It is then congruent that these organisms would optimize transport and reduce replication costs to save energy by streamlining their genomes (Button, 1991, Sowell et. al., 2009). Several of our findings shed light on the ever-expanding knowledge about acidophiles ecology and the acid resistance systems that maintain this proton gradient. Mainly, the increased paralog frequencies in COG categories possibly related to energy production, DNA repair and biofilm formation. The investigation of which functions might be in greater copies in acidophiles is an interesting topic for future research, as it may uncover novel survival mechanisms for acidophiles. Similarly, acid related genes shared between acidophiles could be hidden amongst the proteins without functional annotation.

## Supporting information

Supplementary Fgures

## 6. Author Contributions

DC, GN and DH designed the research. DC performed the research. DC, DH and GN analyzed the data. DC and DH wrote the paper. GC and EV participated in the construction of the final manuscript. All authors read and approved the final manuscript.

## 7. Acknowledgments

DH was supported by Fondecyt 1181717 and Programa de Apoyo a Centros con Financiamiento Basal AFB170004 to Fundación Ciencia & Vida.

