## Supplementary Fgures for "A large-scale genome-based survey of acidophilic Bacteria suggests that genome streamlining is an adaption for life at low pH"

### *Supplementary Figures*

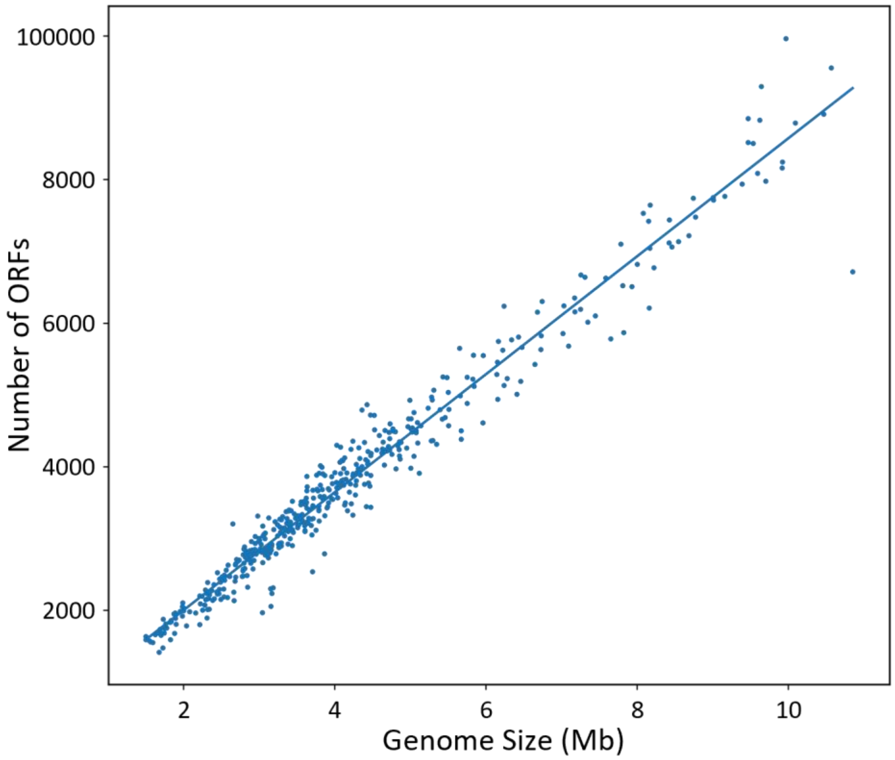


**Supplementary Figure 1.** Number of ORFs versus genome size. Points are the species averages of the number of ORFs and genome size. Pearson's correlation coefficient is 0.98, with a p-value lower than $10^{-320}$.

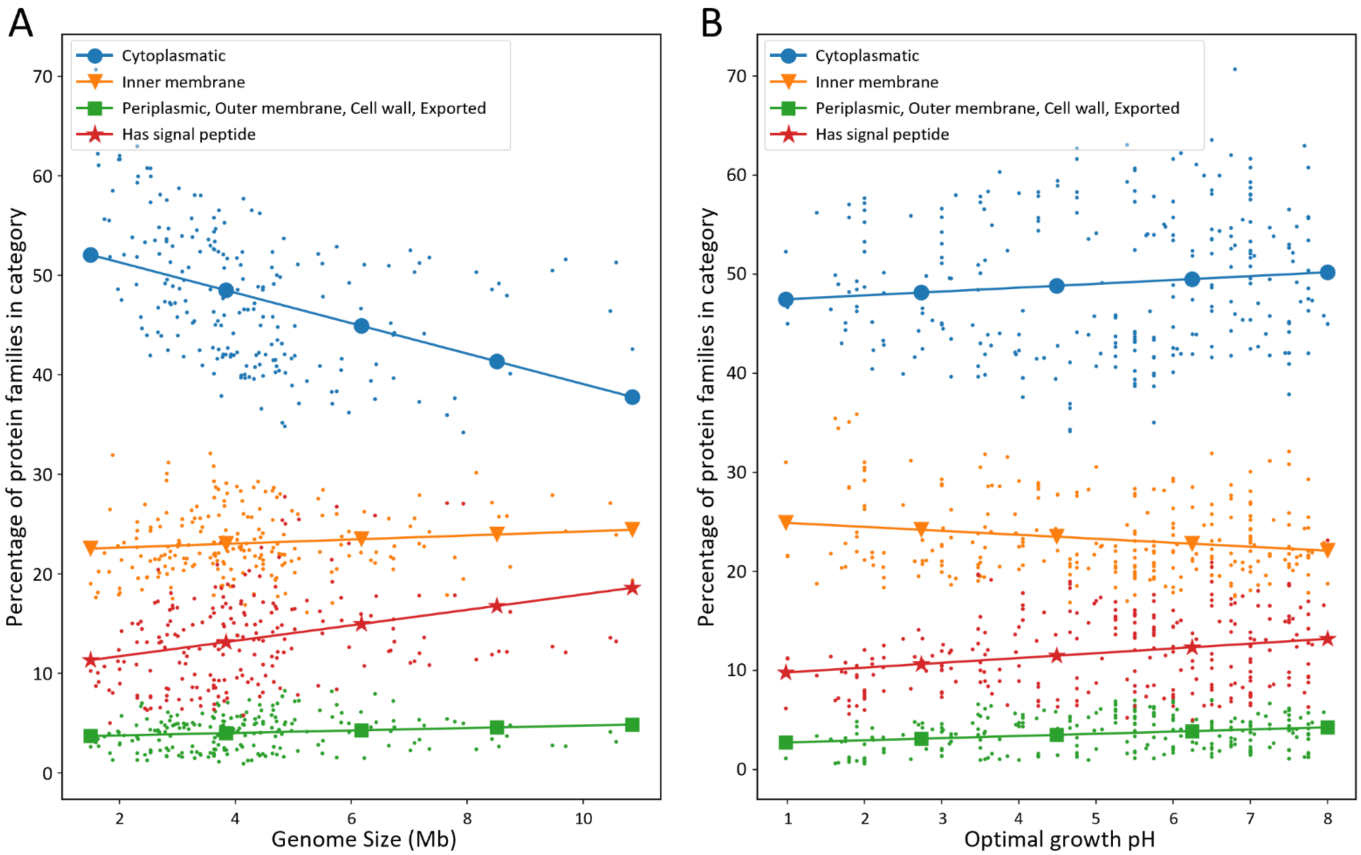


**Supplementary Figure 2.** Disentangling the effects of genome size changes from subcellular localization correlations. **(A)** Subcellular localization vs Genome size. Analog to figure 9 but vs genome size and only for neutrophiles (pH 6-8), to isolate the pH influence. Pearson's correlation coefficient and p-value are respectively -0.4 and $2*10^{-9}$ for cytoplasmic, 0.11 and 0.12 for inner membrane, 0.13 and 0.06 for Periplasmic, Outer membrane, Cell wall and Exported, and 0.3 with $1.3*10^{-5}$ for proteins with a signal peptide. **(B)** Subcellular localization vs pH in small genomes. Analog to figure 9 but for genomes under 4 Mb, where there is no correlation between genome size and pH (p-value = 0.15). Pearson's correlation coefficient and p-value are respectively 0.1 and 0.1 for cytoplasmic, -0.19 and $1.8*10^{-3}$ for inner membrane, 0.24 and $1.3*10^{-4}$ for Periplasmic, Outer membrane, Cell wall and Exported, and 0.24 with $1.0*10^{-4}$ for proteins with a signal peptide.

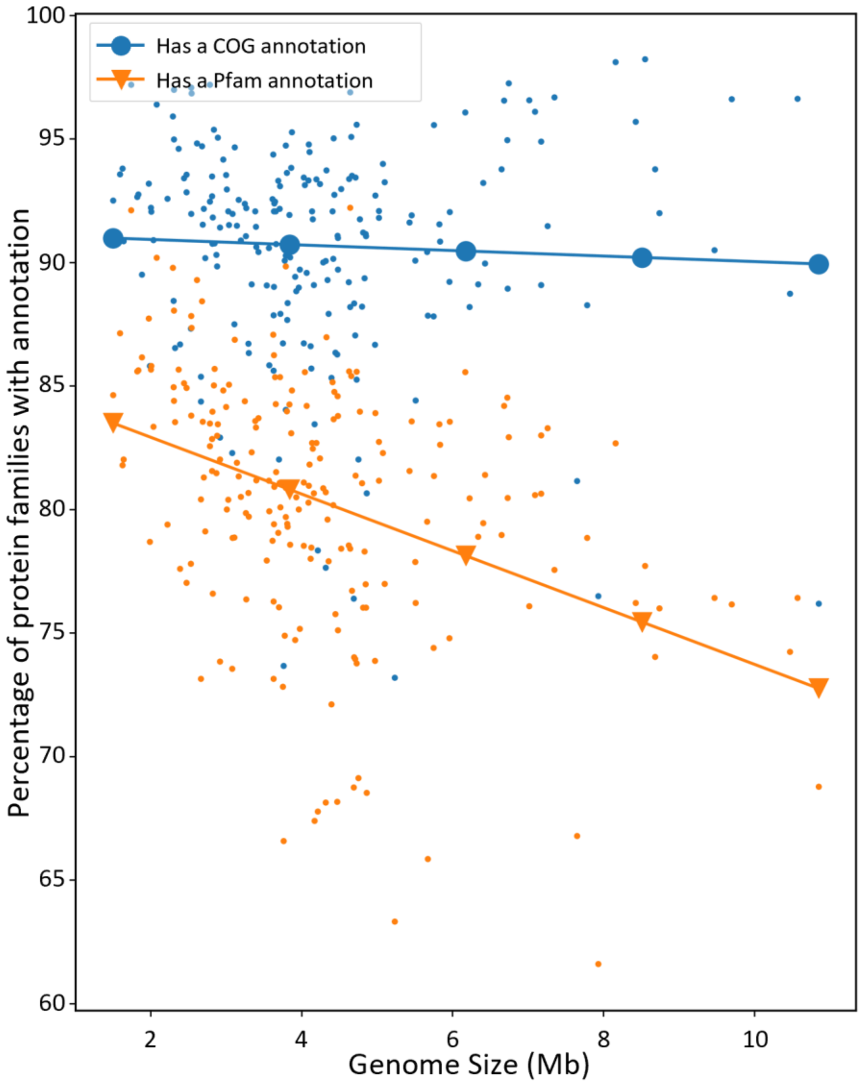


**Supplementary Figure 3.** Percentage of proteins with a functional annotation versus genome size. The percentage of proteins with either COG annotation or Pfam annotation was plotted against mean genome size. Only neutrophiles (Optimal growth pH 6-8) were considered in this analysis to correct for pH influence. Pearson's correlation coefficients are -0.02 and -0.35 with p-values 0.76 and $2.28*10^{-7}$ for COG and Pfam annotations, respectively.

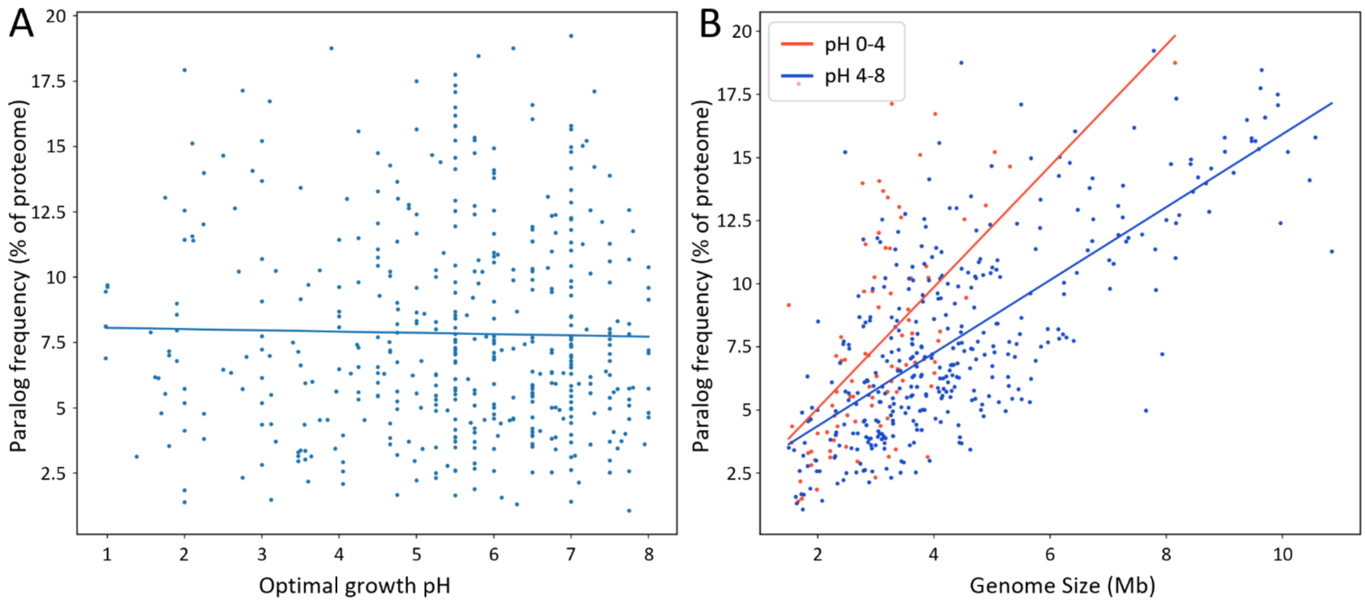


**Supplementary Figure 4.** General paralog frequency tendencies. **(A)** Paralog frequency vs pH. Ortholog groups with more than one protein in the same genome were defined as paralog groups. The percentage of a proteome that belongs in paralog groups (paralog frequency) was plotted against pH. Pearson's correlation coefficient is -0.02, with p-value 0.67. **(B)** Paralog frequency vs genome size at different pH ranges. Pearson's correlation coefficients are, respectively: 0.65 and $2.96*10^{-55}$ for the full range, 0.57 and $4.47*10^{-9}$ for pH 0-4, and 073 with $1.97*10^{-61}$.

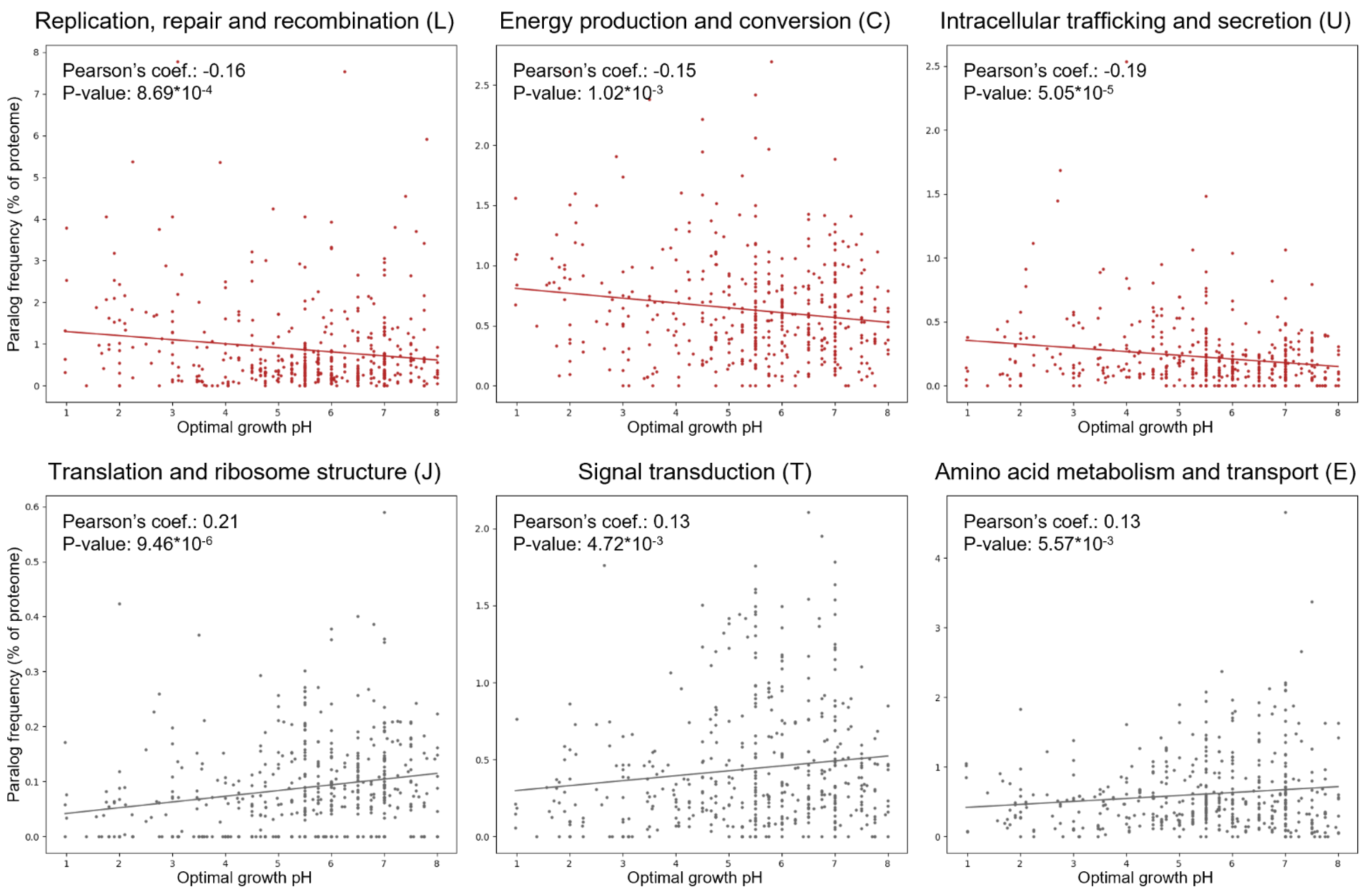


**Supplementary Figure 5.** Paralog frequencies vs pH by COG category plots. Analog to figure 11, the individual scatterplot of paralog frequency vs pH for each COG category with statistically significant correlations (p-value <0.01) are shown. Positive correlations are indicated grey and negative correlations in red. Regression lines are shown, and their projected paralog frequencies at pH 1 and pH 7 were used in figure 11.
